# Co-translational binding of importins to nascent proteins

**DOI:** 10.1101/2022.11.02.514836

**Authors:** Maximilian Seidel, Natalie Romanov, Agnieszka Obarska-Kosinska, Anja Becker, Nayara Trevisan Doimo de Azevedo, Jan Provaznik, Sankarshana R. Nagaraja, Jonathan J. M. Landry, Vladimir Benes, Martin Beck

## Abstract

Various cellular quality control mechanisms support proteostasis. While, ribosome-associated chaperones prevent misfolding of nascent chains during translation, importins were shown to prevent the aggregation of specific cargoes in a post-translational mechanism prior the import into the nucleoplasm. Here, we hypothesized that importins may already bind ribosome-associated cargo in a co-translational manner. We systematically measured the nascent chain association of all importins in *Saccharomyces cerevisiae* by selective ribosome profiling. We identified a subset of importins that bind to a wide range of nascent, often uncharacterized cargoes. This included ribosomal proteins, chromatin remodelers and RNA binding proteins that are aggregation prone in the cytosol. We show that importins act consecutively with other ribosome-associated chaperones. Thus, the nuclear import system is directly intertwined with nascent chain folding and chaperoning.

**One-Sentence Summary:** We describe an unanticipated connection between co-translational protein chaperoning and the nuclear import system.

## Main Text

Faithful protein biogenesis and the maintenance of a functional proteome poses a logistic burden for cells (*1*). Errors in proteostasis result in protein aggregation, consequently leading to pathogenic phenotypes (*2*). Therefore, it is crucial to ensure the quality of nascent proteins already in the vicinity of the ribosome. Nascent proteins are supported by an array of different co-translationally acting quality factors including nascent chain chaperones, nascent chain modifiers and translocation factors such as the signal recognition particle (SRP) and their protein complex partner subunits (*3, 4*). Ultimately, their synergistic action prevents intramolecular misfolding and ensures the reliable formation of stable multidomain arrangements (*3, 4*).

Importins (also called karyopherins) are nuclear transport receptors (NTRs) that bind the nuclear localization sequences (NLSs) of their cargo in the cytoplasm and facilitate its passage through nuclear pore complexes (NPCs) into the nucleoplasm (*5*). Moreover, importins contribute to proteostasis (*6*). *in vitro*, importins inhibit the precipitation of basic, aggregation-prone cargoes by preventing their unspecific interaction with cytosolic polyanions such as RNA (*7*). This has been shown for specific ribosomal proteins and histones (*7*). The importin transportin²2 (Kap²2) suppresses phase separation of RNA-binding proteins such as FUS and interference with cargo binding causes non-native phase transitions (*8–11*). Further, importins disaggregate NLS-bearing cargoes and even rescue neurodegenerative phenotypes *in vivo* (*8*–*11*). Beyond FUS, similar mechanisms may be relevant for TDP-43, TAF15, EWSR1, hnRNPA1, hnRNPA2, arginine-rich proteins and the spindle assembly factor TPX2 (*10, 12, 13*). These previous studies addressed post-translational mechanisms and inferred the molecular binding mode of importins from a limited number of individual cargoes. If importins bind to the nascent chains of their cargo in a co-translational manner, and if so, on which binding sites they generally act, remained unknown.

We reasoned, that during the translation of many proteins that are destined to bind nucleic acids in the nucleus, basic patches are exposed as nascent chains. Protein folding in an RNA-rich environment such as the cytosol may thus critically depend on shielding of the respective patches. We therefore hypothesized that importins may bind to nascent chains. In this study, we systematically measured nascent chain association of all 11 importins in *Saccharomyces cerevisiae*. We used selective ribosome profiling (SeRP) (*14*) to quantify the co-translational binding of importins to nascent proteins in a translatome-wide manner. Our approach led to the identification of a specific subset of importins that co-translationally associate with various cargoes, many of them remained previously unidentified. We show that importins act consecutively with other nascent chain chaperones, in particular on nucleic acid binding proteins. We propose a model in which cargo complex formation is intertwined with nascent chain chaperoning to promote the faithful biogenesis of nuclear proteins. Our findings may have wider implications for our understanding of proteostasis in eukaryotes and provide new perspectives on neurodegenerative disease.

### Selective ribosome profiling identifies the co-translational binding of importins to nascent cargoes

To systematically assess co-translational engagement of importins with nascent cargo (**Fig. 1A**), we used selective ribosome profiling (SeRP) (*14, 15*). This method relies on the affinity purification of co-translational interactors, in this case importins, whereby subsequent sequencing of ribosome-protected mRNA fragments serves as a quantitative proxy for positional chaperone association. It enables the quantification of co-translational binding of nascent chain chaperones to their substrates in a discovery mode for the entire translatome. Furthermore, SeRP systematically unravels the position of binding sites within the relevant open reading frames (ORFs) and thus provides their biophysical properties (*16*–*19*). For affinity purification, we systematically tagged all 11 yeast importins (**Fig. S1A, S1B**) with C-terminal twin-StrepII tags (*20*) using a scar-free cloning technique preserving endogenous 3’ untranslated regions (*21*). We applied the primary amine reactive crosslinker DSP to stabilize potentially transient interactions of importins which may be susceptible to RanGTP throughout lysis (**Figs. S1A, S1C**) as previously described (*15, 16, 22*). After RNase I digestion and enrichment of the ribosome-nascent chain complexes (RNC), we purified the respective co-sedimented importins from the RNCs (**Fig. S1D**). We acquired SeRP data sets by sequencing the ribosome-protected fragments in four biological replicates for each of the 11 importins and a no-bait wildtype strain. We processed the sequencing reads as previously described to obtain the ribosome-protected footprints (**Fig. S1E**) (*14*). The resulting translatome-wide data set captures ribosome footprints for all mRNAs that are affinity-enriched for the respective importins. Pearson correlation between replicates was overall larger than between different conditions (**Fig. S2**). To systematically identify potential hits, we developed a pipeline to short-list candidate profiles that considers both the IP and total translatome relative to a no-bait wildtype control (**Fig. S3**). We used a manually curated list of cargoes of Srp1 and Kap95 from the literature as ground truth (**Fig. S3B** and **Table S1**) and an area under the curve (AUC)-value as a metric to identify co-translational cargoes. We note that our ground truth may contain cargoes that only bind post-translationally, which would result in a conservative, over-estimation of the false discovery rate. Subsequently, we manually inspected all SeRP profiles short-listed by our analysis approach to create a high-confidence list of hits (**Fig. S4**), in which the hits are statistically elevated over the background (**Fig. S5**).

**Figure 1:**
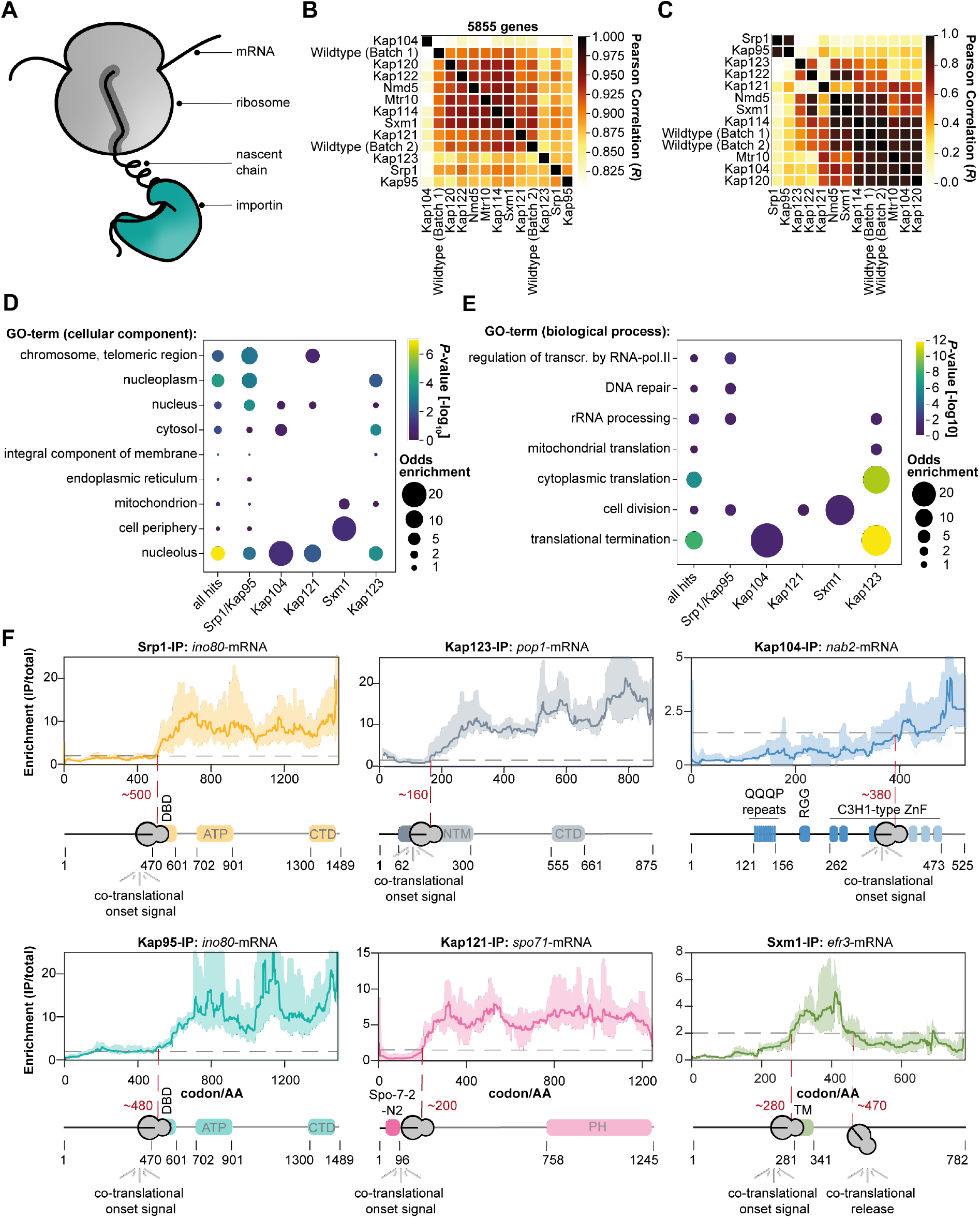
SeRP of importins reveals co-translational binding to cargo. **A**, Scheme illustrating the co-translational binding of importins to a nascent chain. **B**, Pearson correlation of the area under curve (AUC)-values of the selective ribosome enrichment profiles (IP/total) of 5855 genes quantified across the experiments. **C**, Same as **B** but for the 71 manually curated cargoes. **D**, Visualization of the Gene Ontology (GO)-enrichment for cellular compartments. While Srp1-Kap95 enriches chromosomal and telomeric regions, Kap123 shows enrichment for the nucleolus. Only significantly enriched GO-terms are shown (*P*-value < 0.1, not adjusted, Fisher Exact Test relative to all proteins quantified). **E**, Same as **D** but for biological function. While Kap123 enriches for rRNA processing, cytoplasmic translation, and translation termination, Srp1-Kap95 rather enriches for cell division, DNA repair, and transcription regulation. **F**, Representative SeRP profiles. In most cases, importins associate with the nascent chain and subsequently remain bound, whereas Efr3 (Sxm1-SeRP) constitutes an exception. SeRP profiles (IP/total) are shown for the respective mRNA targets from *n*=4 biologically independent replicates (solid lines are averaged across replicates; shades reflect largest to smalls replicate value interval). Grey dashed lines indicate an arbitrary threshold of 2 used for onset estimation (red dashed line). Note, that in the case of *nab2*-mRNA, a threshold of 1.5 was chosen. Domain annotation based on Pfam. transcr.: transcription; RNA-pol. II: RNA-polymerase II; IP: immunoprecipitation; AA: amino acid; DBD: DNA binding domain; ATP: ATP helicase domain; CTD: C-terminal domain; NTM: N-terminal motif; QQQP: glutamine-rich region; RGG: arginine-glycine-glycine domain; C3H1-type ZnF: cysteine-cysteine-cysteine-histidine-type zinc finger domain; PH: Pleckstrin homology domain; TM: transmembrane domain.

Our translatome-wide data, allowed to systematically chart and to compare the co-translational cargo spectra of the different importins. Our approach was very complementary to previous studies that investigated post-translational cargo spectra (*5, 23*–*26*), such that it very accurately identified heterodimeric interactions of importins instead of larger complexes but neglected those that only occur post-translationally. Pearson correlation of the AUC-values for all identified hits suggests a strong separation between the cargoes identified for the individual importins. The data obtained for the beta-type importins Kap114, Kap120, Kap121, Kap122, Mtr10, Sxm1, and Nmd5 largely correlated with negative no-bait controls (**Fig. 1B**). Indeed, very few or no cargoes were detected for this subset. We wondered if this might be related to the detection limit of our method, however, the abundance of importins and the number of identified hits did not correlate (**Fig. S6**). For Srp1, Kap95, Kap123 and Kap121, 28, 27, 30 and 9 co-translationally bound cargoes were detected, respectively. The respective signal was distinct from negative no-bait controls (**Fig. 1C**). In contrast to previous proteomic studies (*27, 28*), we found little overlap between the set of cargoes identified for each importin (**Fig. S7**), with the exception of Srp1 and Kap95 (**see below**). Among the identified hits, gene ontology (GO) analysis revealed enrichment for the nucleus and its sub-compartments underscoring that co-translational binding was specific for nuclear import cargoes (**Fig. 1D**). While cytoplasmic translation and translation termination were enriched in the Kap123-SeRP hit set, regulators of transcription by polymerase II, DNA repair and cell division were found in the Srp1-Kap95 set. We also found that both, Srp1-Kap95 and Kap123 shared enrichment for rRNA processing proteins (**Fig. 1E**). In contrast to Srp1-Kap95 and Kap123 which seemed to be distinct in their function, Kap121 was associated to processes within the nucleolus as well as the chromosome and telomeric regions. Taken together, this data pointed to a model in which Srp1-Kap95, Kap123, and Kap121 prominently associate with a specific set of cargoes in a co-translational manner, while other importins may preferably act post-translationally.

### Co-translational association of importins is enduring

Selective ribosome profiles allow visualization of importin binding events within an open reading frame, as shown for representative examples in **Fig. 1F**. In contrast to profiles previously obtained for the ribosome-associated Hsp70 chaperones Ssb1 and Ssb2 (Ssb1/2) (*16, 17*), chaperonin TRiC (*16*) or the signal recognition particle (SRP) (*18, 19*), the importin-derived profiles suggested that once importin is bound to the nascent protein, it remained tethered (**Fig. 1F**). This is reminiscent of the co-translational interactions previously observed during protein complex formation (*29, 30*). This particular binding pattern may be due to the requirement of RanGTP for the dissociation of import complexes that is absent in the cytosol (*31, 32*). Thus, importins constitute an enduring chaperoning system that holds onto its substrates from synthesis in the cytosol until release into the native context in the nucleus. A notable exception was Efr3 (**Fig. 1F**), which is annotated as a plasma membrane protein. Nevertheless, a strong signal was observed for the importin Sxm1 that binds to the nascent chain of Efr3 approximately at codon 280. Interestingly, it was released ∼180 codons downstream, similar to the transient binding mechanisms of ubiquitous co-translational chaperones and the SRP.

### Srp1 and Kap95 mutually bind to nascent cargo

Srp1 (importin-alpha) and Kap95 (importin-beta) represent the classical nuclear import pathway in yeast (*33*). In contrast to other beta-type importins that directly bind the NLSs of their cargoes, this classical pathway requires importin-alpha as an adaptor. The interaction with importin-beta liberates the autoinhibitory NLS of importin-alpha from the NLS binding groove. This structural rearrangement results in the activation of the Srp1-Kap95 heterodimer that in turn binds to the cargo NLS (*34*). To explore if Srp1 and Kap95 mutually bind to nascent chains, we compared the AUC-fold change of the Srp1-to the Kap95-SeRP experiments, which was highly correlated (**Fig. 2A**). 27 cargoes are common to both experiments (**Figs. 2A, 2B**). In addition, nascent Srp1 itself was bound by Kap95 reflecting a co-translational protein complex formation (**Figs. 2B, 2C**). The respective SeRP profile showed an onset approximately at codon ∼150 (**Fig. 2C**), suggesting the interaction with Kap95 occurred once the synthesis of the importin-beta binding domain (IBB) was completed.

**Figure 2:**
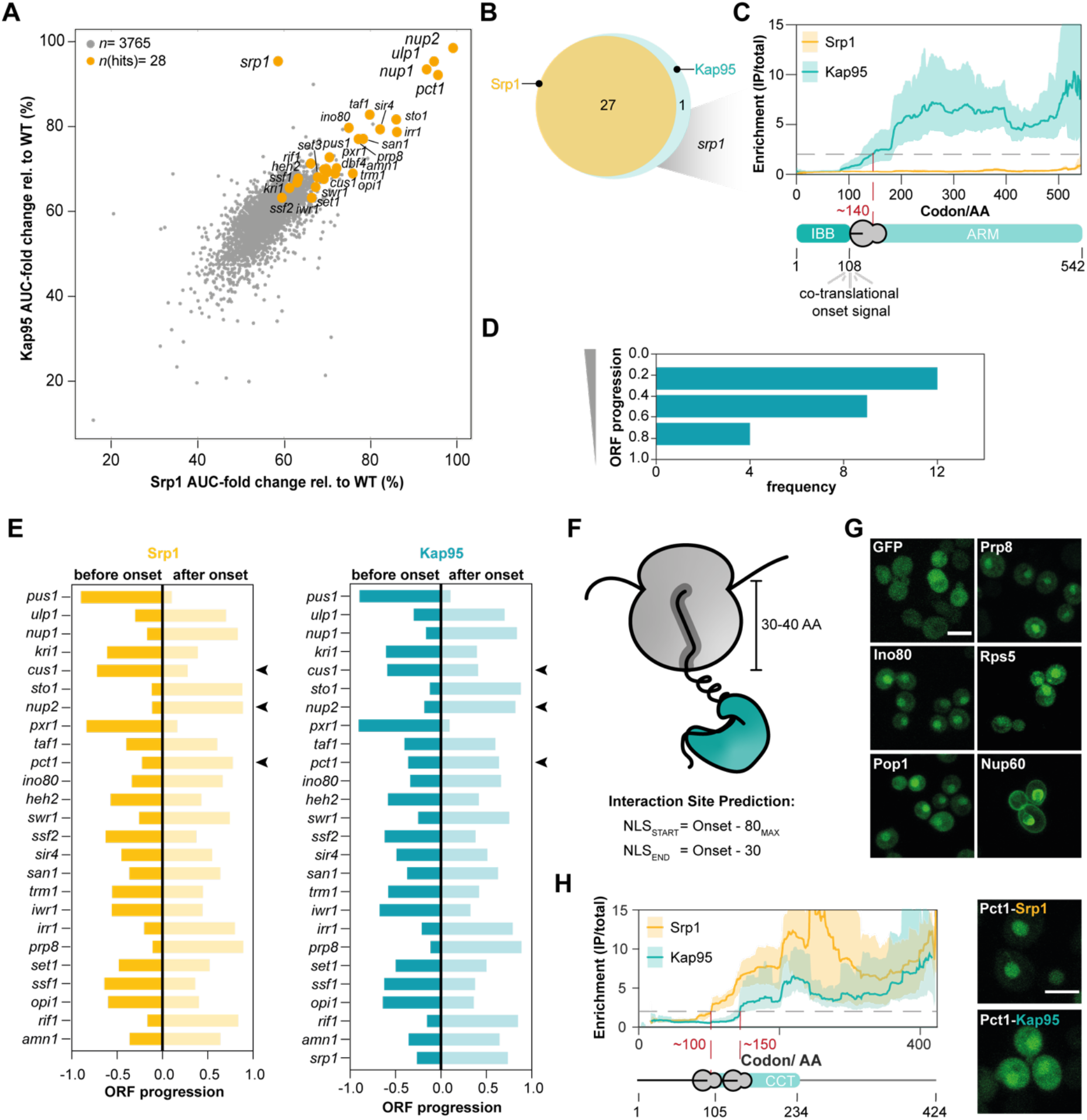
Srp1 and Kap95 synchronously bind to nascent cargoes. **A**, Scatter plot of the AUC-values of genes quantified in Srp1 in comparison to Kap95 (detailed in **Materials and Methods**). The identified hits are highlighted in orange. **B**, Venn diagram showing the overlap of the identified cargoes. **C**, Enrichment profile of *srp1-*mRNA in Srp1- and Kap95-SeRP experiments. Kap95 binds to nascent Srp1 at codon ∼140 corresponding to the release of the entire IBB domain. SeRP profiles (IP/total) are shown for the respective mRNA targets from *n*=4 biologically independent replicates (solid lines are averaged across replicates; shades reflect largest to smalls replicate value interval). Grey dashed lines indicate an arbitrary threshold of 2 used for onset estimation (red dashed line). **D**, Distribution of onsets shows an N-terminal preference. **E**, Comparison of the onsets observed in the Srp1- and Kap95-experiments. Arrowheads indicate 3 slightly divergent cases. **F-G**, Peptides upstream of the observed onsets are sufficient for nuclear localization of the respective GFP fusion proteins. **G**, Representative confocal images for GFP fusions with peptides from the indicated cargoes. Scale bar: 5 µm. **H**, SeRP profiles as in **C** but for *pct1* indicate a slightly shifted onset of Srp1 and Kap95; the N-terminally localized peptide shows stronger nuclear localization apparent in confocal slices (as in **F**). IP: immunoprecipitation; AA: amino acid; IBB: importin-beta binding domain; ARM: armadillo repeat; NLS: nuclear localization sequence; GFP: green fluorescent protein; CCT: choline-phosphate cytidylyltransferase.

We therefore wondered if the onset observed for the Srp1 and Kap95 association occurred at similar positions within the relevant ORFs. SeRP profiles for both, Srp1 and Kap95 showed a pronounced N-terminal preference, contrasting other co-translationally acting importins (**Fig. 2D** and **Fig. S8**). On the level of individual ORFs, simultaneous binding of Srp1 and Kap95 was observed (**Fig. 2E** and **Fig. S9**), with very few exceptions, suggesting heterodimer formation prior to cargo binding. This finding suggested that the above-introduced mechanism of Srp1-Kap95 heterodimer activation, which has been elucidated by structural and biochemical analysis for a smaller set of substrates, appears to be broadly applicable to the co-translational formation of cargo complexes (*25, 34*).

To address the functional relevance of the observed onsets, previous studies used genetic perturbation and biochemical assays (*16, 18, 19, 22, 29, 30*). We queried whether the respective peptides upstream of the onset would be sufficient for nuclear localization. Therefore, we generated GFP fusions of peptides within a 40 amino acid sequence window upstream of the onset, accounting for the emergence of the peptide from the ribosomal exit tunnel (**Fig. 2F**). While GFP without any fusion peptide was present throughout the entire cell, peptide fusions of 5 randomly selected cargoes (Srp1-Kap95: Ino80, Prp8; Kap123: Rps5, Pop1, Nup60) were sufficient for nuclear localization (**Fig. 2G**). In case of Pct1, for which a slightly shifted onset of Kap95 with respect to Srp1 was observed, we found that the N-terminally localized peptide showed a stronger nuclear enrichment (**Fig. 2H**).

### Prediction of classical NLSs in Srp1-Kap95 cargoes

To map putative cNLS in the proteins identified as hits, we ran AlphaFold-Multimer (*35*) structure prediction for pairs of Srp1 and consecutive overlapping fragments of the respective protein sequences (see **Materials and Methods**). For all hits, we obtained at least one prediction with a fragment occupying the NLS binding site of the Srp1. Some hits contained two or more NLSs predicted with similar scores. The predicted NLSs frequently occurred at the N-terminal region in agreement with the N-terminal preference found within the SeRP data (**Fig. 2D**). All known NLS motifs were predicted with top scores (**Table S2** and **S3**) validating our procedure. The structural superposition (**Fig. 3A**) and structure-based sequence alignment revealed that most of the predicted motifs exhibit sequences resembling classical NLS (cNLS) motifs of Srp1 of either the monopartite (K-K/R-X-K/R) or bipartite (K/R-K/R-X_10-12_-K/R_3/5_) type (**Fig. 3B**) (*36*), whereby the linker region can be considerably longer. Some sequences, however, were very different from the sequence consensus or bound to the NLS binding site in the opposite direction and might correspond to false positive predictions or non-canonical NLSs. Altogether, these results confirmed that the identified target proteins bind to Srp1 and allow a prediction of the corresponding cNLS motifs.

**Figure 3:**
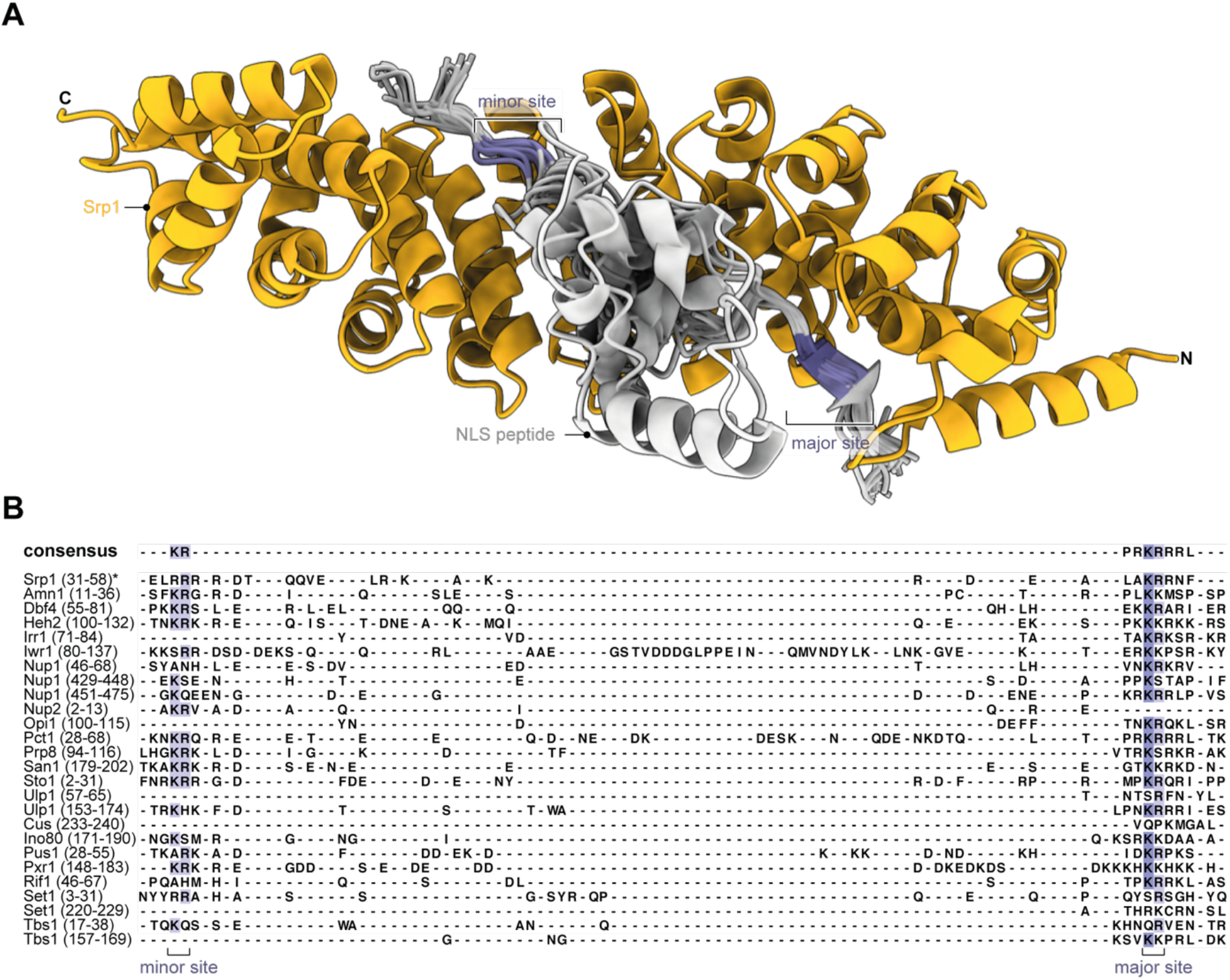
Co-translationally bound Srp1 cargoes have predicted cNLSs. **A**, Srp1 (AA 70-512) structure (orange) with peptides modeled using AlphaFold-Multimer. Only predictions i) with the ipTM+pTM score > 0.7, ii) bound to the canonical NLS-binding site in Srp1, iii) bound in the N-to C-terminus orientation as known from crystal structures of Srp1-NLS are shown (PDB: 1WA5) (*37*). The entire data set is listed (**Table S3**). **B**, Structure-based sequence alignment derived from subpanel **A** display hallmarks of cNLSs. Regions of the target sequences are shown in the alignment indicated in parentheses (in AA). Peptides are colored by sequence conservation. *inhibitory peptide of Srp1 (PDB: 1WA5) (*37*).

### Kap123 cargoes act in early stages of ribosome biogenesis

Ribosomal proteins (r-proteins) are synthesized in the cytoplasm and are transported into the nucleolus where they associate with ribosomal RNAs (rRNA). Out of the 87 yeast r-proteins, 13 were detected in our screen, including 5 paralogous pairs. 12 out of 13 detected r-proteins co-translationally engaged with Kap123, the other one with Kap104 (**Fig. S4**). We found that for the paralogous r-proteins, the selective ribosome profiles were identical in shape, and enrichments varied according to their paralog-specific expression levels (*38*) (**Fig. S10A**). Moreover, none of the identified r-proteins overlapped with the substrate spectrum of known r-protein chaperones (*39*) (**Fig. 4A**), suggesting a unique functional role of Kap123 in ribosome biogenesis. Specifically, the identified r-proteins suggested that Kap123 is relevant for early stages of 60S and 90S ribosome biogenesis (**Fig. 4B**). This notion is further supported by other identified cargoes that include several early ribosome biogenesis factors such as e.g. Nug1, Noc2, Ecm16 (**Fig. 1E, Fig. S4**) (*40*–*43*).

**Figure 4:**
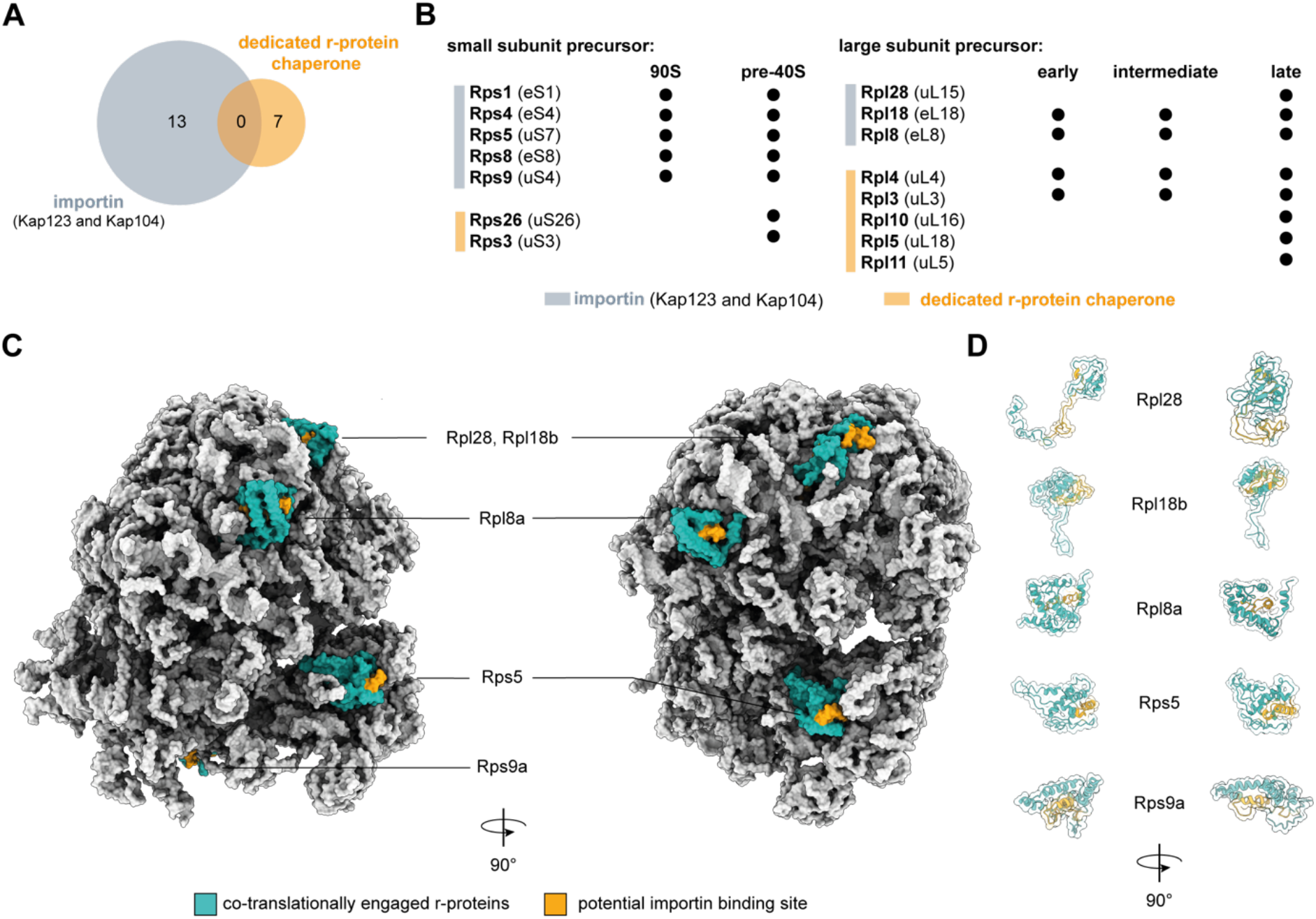
Engagement of Kap123 with nascent chains reveals novel aspects in ribosome biogenesis. **A**, Venn-Diagram of cargo overlap between importins and dedicated r-protein chaperones (Tsr2, Yar1, Acl4, Rrb1, Sqt1 and Syo1). The importins Kap123 and Kap104 chaperone unique r-proteins which else would not be covered by the substrate spectrum of r-protein chaperones. **B**, Importin chaperoned r-proteins are required in early ribosome biogenesis. **C**, Potential Kap123 and Kap104 binding moieties are buried within mature 80S ribosome. Co-translationally engaged r-proteins are highlighted in turquoise with their potential importin binding site highlighted in orange. Peptides highlighted in orange represent -50 to -30 of the onsets. **D**, same as for **C**, but represented as cartoons. Depicted structures can be accessed at the PDB: 4V7R (*44*).

Interestingly, when we depicted the apparent onsets for the interaction of Kap123 into a mature 80S ribosome structure (PDB: 4V7R) (*44*) we noticed that they typically mapped to the C-terminally located structured domains of the respective r-proteins (**Figs. 4C, 4D** and **S8**). Within the mature 80S ribosome, the respective sites were engaged in various contacts suggesting an inaccessibility for importin binding. A possible interpretation of this observation is that faithful structural rearrangements of r-proteins during biogenesis ultimately renders ribosomes invisible to the nuclear import system to prevent nuclear re-import.

### Ssb1/2 chaperoning occurs upstream of importin binding

Upstream of their structured domains that are bound by Kap123, the respective r-proteins typically contain a charged N-terminal patch that appears intrinsically disordered in their primary structure, but is engaged in contacts with rRNA in the mature ribosome. We therefore wondered whether the broadly acting nascent chain chaperones Ssb1/2 (*17*) that are known to bind intrinsically disordered, charged patches, could act upstream of Kap123. We therefore systematically analyzed the co-occupation of importin chaperoned nascent chains with Ssb1/2, and the Hsp60 TRiC/chaperonin using previously published data sets (*16, 17*) (**Fig. S10B**). We found that the nascent chains of all r-proteins bound by Kap123 and Kap104 are also captured by Ssb1/2, and in the case of Rpl8a, Rps1a/b, Rps5 and Rps9b also by TRiC. Across the Ssb1 co-chaperoned r-protein SeRP profiles, we found that Ssb1 binding temporarily preceded Kap123 binding (**Figs. 5A, 5B**, and **Fig. S10C**). A notable exception was Rpl28 that has not been reported as a substrate for other nascent chain chaperones and may not require Ssb1/2-binding due to its unstructured N-terminus captured by Kap123 (**Fig. 5C**). Previous mass spectrometry data indicated that a subset of the yeast proteome is rendered aggregation-prone in absence of Ssb1/2 (*45*). While Ssb1/2 substrates were depleted in nascent Srp1-Kap95 or Kap121 cargoes as compared to the nuclear proteome, they were enriched in the corresponding nascent Kap123 cargoes (**Fig. S10D**). Interestingly, nascent Kap123 cargoes very frequently aggregated in the absence of Ssb1/2 (**Fig. S10D**), suggesting that both processes are intertwined. Taken together, this analysis suggested a Ssb1/2-importin handover mechanism for substrate recognition (**Fig. S10E**). Since Ssb1/2 is thought to retain hydrophobic and positively charged nascent peptides in a linear, degenerated form (*16, 17*) it may facilitate co-translational importin cargo recognition.

**Figure 5:**
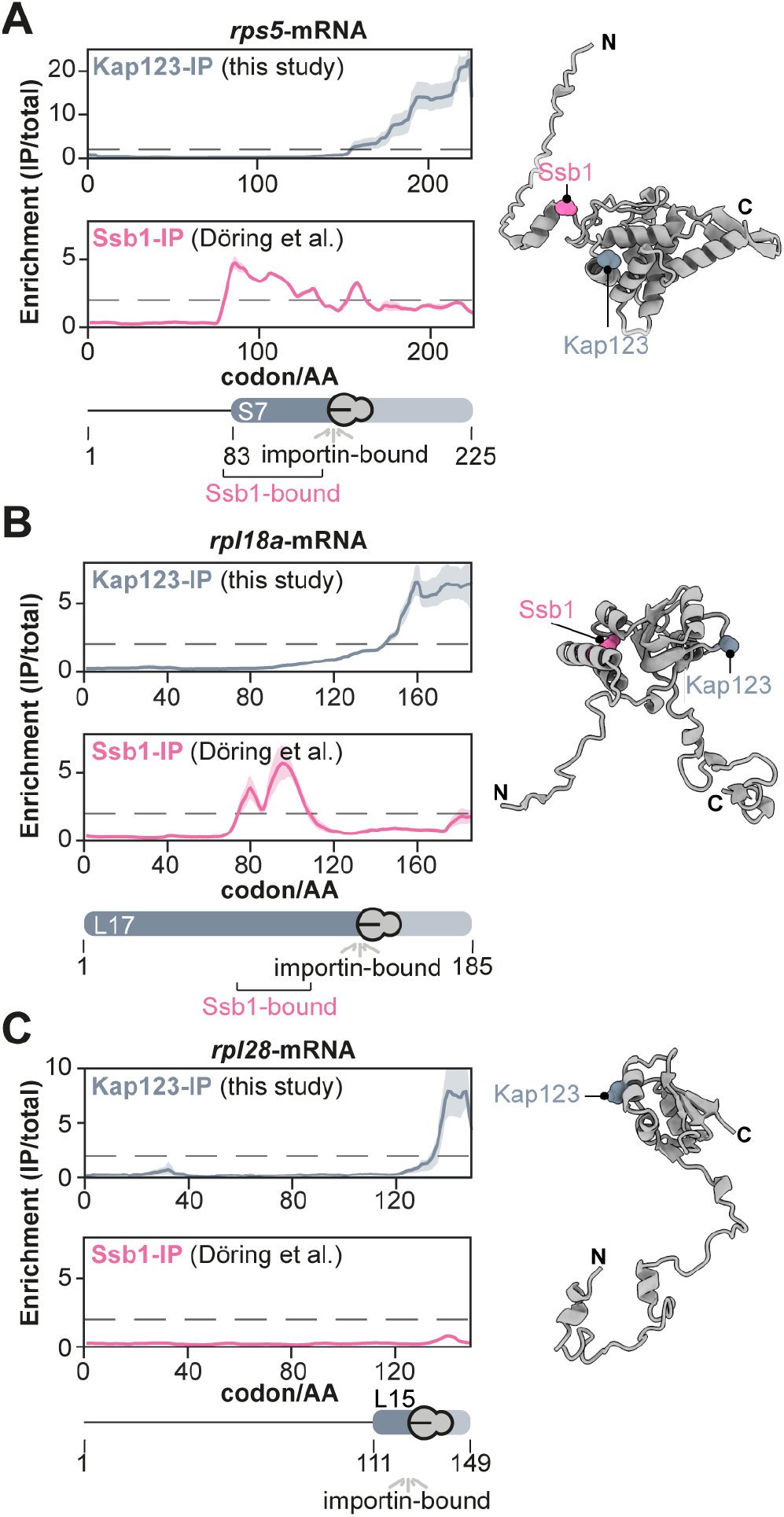
Nascent chaperoning and cargo complex formation of r-proteins. SeRP profiles and AlphaFold models of **A**, Rps5, **B**, Rpl18a, and **C**, Rpl28. **A**, and **B**, The SeRP profiles reveal, that for nascent Rps5 and Rpl18a, Ssb1 first binds to the nascent chain. Subsequently, it is released prior to Kap123 binding. **C**, In contrast, Rpl28 does not require Ssb1 chaperoning. Nevertheless, Kap123 binds C-terminally. The folds exemplify the structural organization of many r-proteins. A nascent r-protein recognized by Kap123 contains an N-terminally disordered, charged patch and a consecutive structured domain. SeRP profiles (IP/total) are shown for the respective mRNA targets from *n*=4 biologically independent replicates (solid lines are averaged across replicates; shades reflect largest to smalls replicate value interval). Grey dashed lines indicate an arbitrary threshold of 2 used for onset estimation. SeRP for Ssb1 (*17*) represents profiles generated from *n*=2 biologically independent replicates. AlphaFold models of r-proteins were obtained from the AlphaFold database (https://alphafold.ebi.ac.uk/) (*46, 47*). C-terminal binding sites (40 AA downstream of onset) are labeled for Ssb1 and Kap123 within the structures, respectively. IP: immunoprecipitation; AA: amino acid.

### Co-translational importin binding sites have distinct biophysical properties

While small molecules can diffuse across NPCs, larger molecules above ∼30-50 kDa require active transport (*48*). We therefore asked if co-translationally associated cargoes were above this size threshold. Our analysis showed that the majority of the Srp1-Kap95 chaperoned cargoes exceed the size threshold and thus depend on active import. This was less pronounced for the Kap121 cargoes and stood in strong contrast to the Kap123 cargoes. In the latter set, small proteins with a median size of only ∼30 kDa were particularly enriched suggesting that the majority of which could also passively enter the nucleus if they were not bound to Kap123 (**Fig. S11A**). Concomitantly, many of the Kap123 cargoes were highly abundant and synthesized with an exceptionally high translation rate (**Figs. S11B, S11C**). These findings point to a model in which co-translational cargo complex formation may mask potentially harmful biophysical properties, in particular of Kap123 cargoes.

Gene Ontology analysis indicated that many co-translational cargoes encode for ribonucleic acid binders (**Fig. 6A**). The apparent onsets of importins within the ORFs of co-translational cargoes were enriched for specific types of domains, namely tRNA methyltransferases as well as DNA-, RNA-or histone processing domains. To a much lesser extent, they occurred at sites of protein-protein interactions or membrane association (**Fig. 6B**). Since nucleic acid binding domains are often charged, we assessed the isoelectric point (pI) of the proteins detected by our screen. In comparison to the entire nuclear proteome, co-translationally bound cargoes had higher pI-values and were enriched for lysine and arginine, in particular Kap123 cargoes (**Fig. 6C, Fig. S11D**). Interestingly, co-translational cargoes of importins bear strong positive charges as compared to the substrates of the ubiquitous chaperones Ssb1/2 and TRiC (**Fig. S11E**), stressing the unique role of importins. The local amino acid signature upstream of the observed onsets for Srp1 and Kap123 displays a strong enrichment for the positively charged residues lysine and arginine as compared to the entire protein (**Figs. 6D**). At last, we analyzed if the elongation fidelity is affected by the compositional bias at importin binding sites. We noticed an increased ribosomal occupancy upstream to the observed onset suggesting a decrease in elongation prior to importin binding (**Figs. 6E, 6F**). Interestingly, the charged lysine and arginine residues that are enriched at the observed onsets of many cargoes, are not only a hallmark of the NLS motif but they are also associated with less abundant tRNAs (*49*) and thus likely reduce translation speed to warrant importin binding. These findings generalize the basic chaperoning function of importins for numerous cargoes and provides more detailed insights about the chaperoned cargo sites and their biophysical properties.

**Figure 6:**
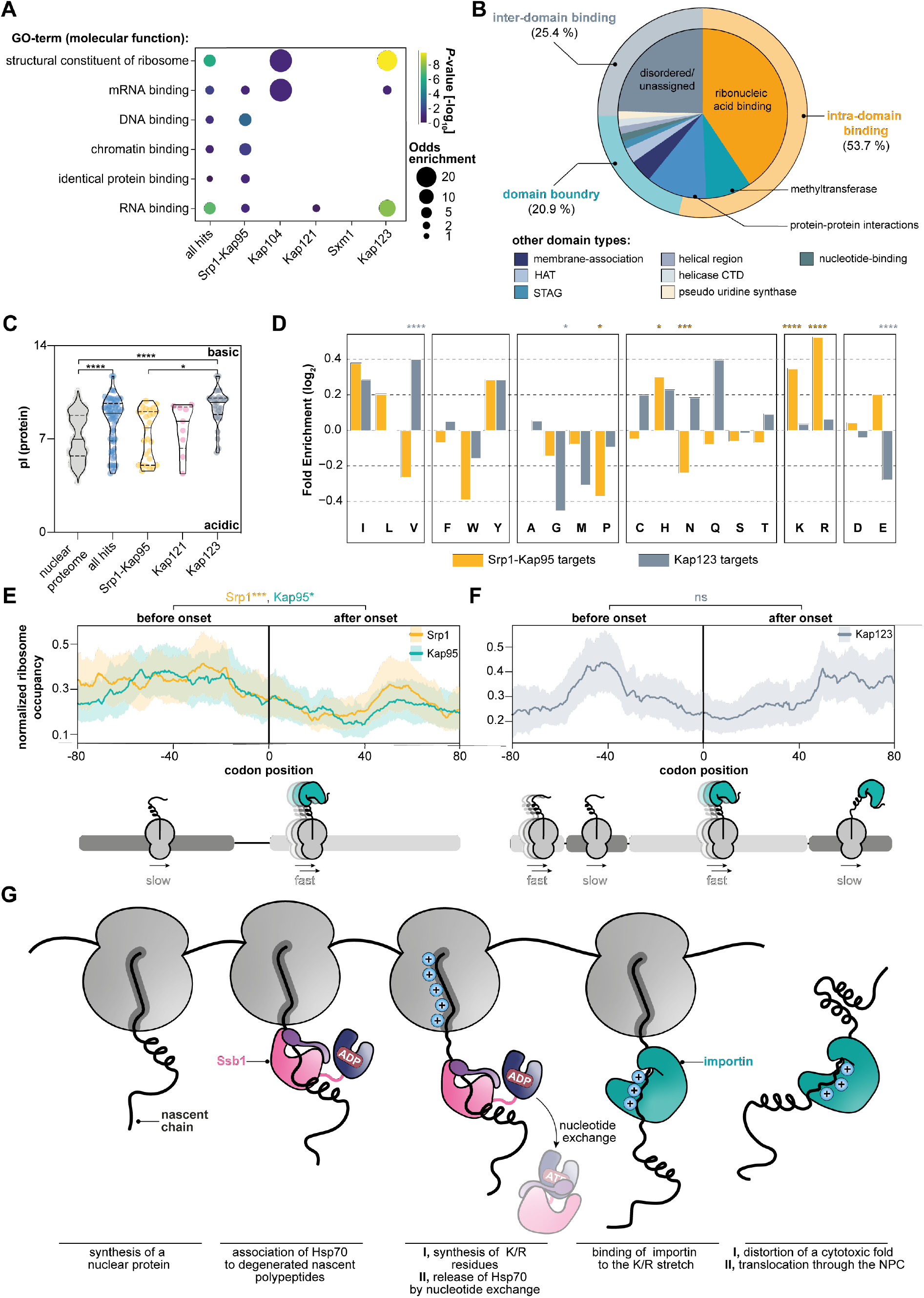
Importins protect positively charged ribonucleic acid binding domains. **A**, Visualization of the Gene Ontology (GO)-enrichment for molecular functions. Co-translationally targeted cargo shows strong enrichment for DNA-, RNA-, chromatin-, protein-binding properties, and structural proteins of the ribosome. Only significantly enriched GO-terms are shown (P-value < 0.1, not adjusted, Fisher Exact Test relative to all proteins quantified). **B**, Onsets are frequently observed at ribonucleic acid binding sites. Apparent onsets were mapped as described in **Materials and Methods** and classified according to their annotated function. **C**, Importins capture preferentially positively charged nascent cargoes. While the pI-values of nuclear proteins are distributed bimodally, nascent cargoes are shifted towards high pI-values. Violin plot shows median and quartiles. \*\*\*\**P*< 0.00074 (nuclear proteome, all hits); \*\*\*\**P* <0.001 (nuclear proteome, Kap123); \**P*= 0.032 (Srp1/Kap95, Kap123). Non-parametric Mann-Whitney U-test. **D**, Amino acid enrichment in apparent onsets for nascent Srp1-Kap95 and Kap123 cargoes as compared to the full-length proteins (Fisher Exact Test). \**P*= 0.0101 (Srp1-Kap95; proline); \**P*= 0.0152 (Srp1-Kap95; histidine); \*\*\**P*= 0.0015 (Srp1-Kap95; asparagine); \*\*\*\**P*< 0.001 (Srp1-Kap95; lysine); \*\*\*\**P*< 0.001 (Srp1-Kap95; arginine); \*\*\*\**P*< 0.001 (Kap123; valine). \**P*= 0.0489 (Kap123; glycine). \*\*\*\**P*< 0.001 (Kap123; glutamate). **E**, and **F**, Metagene analysis of the ribosome occupancy at apparent onset sites for nascent Srp1-Kap95 (**E**) and Kap123 (**F**) cargoes. Prior to the onset, an increased occupancy is observed, implying a decrease in elongation. \*\*\**P*= 0.0028 (Srp1: before, after onset); \**P*= 0.0113 (Kap95: before, after onset); ns= 0.8642 (Kap123: before and after onset). Solid lines represent averaged profiles; shades reflect a 95% confidence interval. For **C**,-**F**, ns *P*> 0.05, \**P*< 0.05, \*\**P*< 0.01, \*\*\**P*< 0.005, \*\*\*\**P*< 0.001. **G**, Conceptual model of nascent cargo complex formation. As the nascent chain emerges from the ribosome, it may be bound by ubiquitous chaperones (e.g. Ssb1) that temporarily chaperone structured patches. Once released, importins nascently form cargo complexes and shield charged patches. GO: gene ontology; HAT: histone acetyltransferase; CTD: C-terminal domain; pI: isoelectric point.

## Conclusion

Taken together, our data point to the following model (**Fig. 6G**): Importins bind to the nascent chain of many cargoes during protein synthesis. This mechanism is particularly prominent for basic nuclear and r-proteins that are primarily substrates for Srp1-Kap95, Kap121, and Kap123. Prior to co-translational engagement of importins, many lysins and arginines, which are often constituents of nucleic acid binding domains, are synthesized causing an intrinsic reduction of the translational speed due to their rare codon usage. This decrease in translational fidelity may be beneficial for faithful importin association. In some cases, we even observe a direct handover between temporarily bound Ssb1/2 and importins. This handover might be necessary for cargo recognition by importin in particular for proteins whose importin binding sites become inaccessible in the ternary structure.

Predicted and experimentally characterized NLSs indicate that in some cases, the observed onset may be shifted downstream to the physical importin binding site. This may be explained by the modulation of the availability of the binding site by other nascent chain binders as exemplified for Hsp70 or temporary acetylation of N-terminal NLS by N^±^-terminal acetyl transferases (*50, 51*). Alternatively, domain recognition of importins may explain this phenomenon as exemplified by a recent structural study (*52*).

We propose that the co-translational nuclear import complex formation shields positively charged patches early during biogenesis, in an RNA-rich environment. This may be particularly relevant for nucleic acid binding domains that otherwise may be aggregation-prone in the cytosol. These insights strengthen the notion that importins are part of a basic client chaperone network. It was previously shown that importin 4 (yeast: Kap123) chaperones the basic RPS3A (yeast: Rps1a) (*7*) that otherwise aggregates in the presence of tRNA. Our study highlights that this protection of Rps1a is already established co-translationally. This chaperoning presumably lasts until the nuclear entry of the import complex. In the nucleoplasm, it becomes exposed to RanGTP and the cargo is released into the destined biophysical environment to engage with native interaction partners. Some of the cargo complexes may rely on alternative release cues, such as the histone dimer H2A•H2B, the SUMO-deconjugating enzyme Ulp1, the mRNA binding protein Nab2, Nab4, and Npl3, or the ribosomal protein eS26 (*53*–*57*).

Beyond the co-translational proteostatic function of importins, our study extends the known spectrum of cargoes chaperoned by importins beyond ribosomal proteins, histones, and some RNA binding proteins (*7, 10*). We found 71 unique co-translational substrates (summarized in **Fig. S4**), many of them accounting for previously undescribed nuclear transport cargoes. Surprisingly, histones and many of the r-proteins did not enrich co-translationally. This may be due to the action of additional and very specialized chaperone networks that may protect them from misfolding in a co-translational fashion (*39, 58*). Although importins have been shown to be partially redundant in function and their cargo spectrum (*27, 33*), our data indicate a rather low redundancy in the co-translational binding capacity (**Figs. 1B, 1C** and **Fig. S7**). We find that the Srp1-Kap95 heterodimer, Kap123, Kap121 and to lesser extent Kap104, Sxm1, Mtr10 and Kap122 co-translational act on nascent chains, while Kap114, Kap120, and Nmd5 did not show any significant binding under the conditions tested (**Figs. 1B, 1C**). One may speculate that the subset of importins identified in this study act as co-translational chaperones under optimal growth conditions. Other importins may be required under permissive conditions, for example, Kap114 that is indispensable under saline stress (*59*).

Overall, our findings suggest a role of importins as proteostatic safeguards for nascent nuclear proteins but also open up novel perspectives on previous findings that associated importins with biomolecular condensation. For example, Kap123 and some of the here identified co-translational cargoes (e.g. Nug1 and Noc2) were reported to phase separate upon heat shock (*60*). We speculate that recruitment of Kap123 may ensure reversibility by protecting the RNA-binding patches of cargoes in the respective RNA containing granules. Furthermore, importin-alpha, importin-beta, and the Kap121 homolog importin-²3 were found to co-translationally associate with Nup358-granules that manufacture NPCs in early fly development (*61*). Most importantly, importins were attributed to counteract neurodegenerative disease by enhancing the solubility of nuclear proteins associated with pathological features such as FUS and TDP-43 (*6*). Some of the genes identified in *S. cerevisiae* in our study are known to drive neurodegeneration in humans. Among these genes are *prp8* that is associated with retina pigmentosa (*62*), *efr3* that is mutated in autism spectrum disorder (*63*), and *taf1* that if mutated can cause intellectual disability (*64*). Our study demonstrates that the solubility of such proteins may be enhanced co-translationally to prevent the exposure of aggregation-prone ribonucleic acid binders prior to their full accessibility to the cytoplasm.

## Acknowledgment

We thank Patrick Hoffmann, Filipa Pereira, and Gerhard Hummer for insightful discussions; Erin Schuman, and Alessandro Ori for critical reading of the manuscript. We would like to thank Dingquan Yu and Jan Kosinski from CSSB & EMBL Hamburg for support with AlphaFold modeling. The authors would like to acknowledge the support of the Imaging Facility at the Max Planck Institute of Brain Research. Additionally, the authors like to thank Georg Stoecklin, Lars Steinmetz, and Britta Brügger for the critical assessment of the project. M.B. acknowledges funding by the Max Planck Society and the European Research Council (724349-ComplexAssembly).

## Authors Contribution

M.S. conceived the project, designed experiments, performed experiments, analyzed data, and wrote the manuscript. N.R. developed the analysis pipeline, analyzed data, and wrote the manuscript. A.O.-K. conducted AlphaFold modeling, analyzed data, and wrote the manuscript. A.B. performed experiments. J.J.M.L., N.T.D.d.A, and J.P. performed experiments and analyzed data. S.R.N. performed experiments. V.B. designed experiments and supervised the project. M.B. conceived the project, designed experiments, analyzed data, supervised the project, and wrote the manuscript.

