## Supplementary_Information for "Co-translational binding of importins to nascent proteins"

**This PDF file includes:**

Materials and Methods
Figs. S1 to S11
Tables S1 to S4
Caption for Table S3

**Other Supplementary Materials for this manuscript include the following:**

Table S3

### Materials and Methods

#### Yeast strain design

Scar-free C-terminally twin-StrepII-tagged importin strains were obtained by homologous recombination using the MX4 blaster cassette (*21*). For this method, the MX4 blaster cassette was amplified with gene-specific overhangs to recombine immediately after the endogenous STOP-codon of the gene of interest. These PCR products were then transformed into BY4741 and selected on YPD-high phosphate plates containing 300 µg/mL hygromycin B (ForMedium) and 3 g/L potassium dihydrophosphate (monobasic). In the second round of transformations, MX4 blaster cassette was substituted with a codon-optimized StrepII-tag with flanking gene-specific overhangs. For this transformation, the cells were grown in low phosphate YPD (*21*) and then transformed. Clones were selected on YP-galactose plates. We validated twin-StrepII-tag insertion by PCR.

For the NLS-GFP constructs, we obtained plasmids by Gibson Assembly (NEB) of the NLS flanked with 20 bp overhangs and a pRS316 containing a *tef1*-promoter :: GFP :: *cyc1*-terminator. 5 ng of plasmid was transformed into BY4741 and the cells were selected on uracil drop out plates (ForMedium). All yeast strains generated in this study are listed in **Table S4**.

#### Selective ribosome profiling

Selective ribosome profiling was conducted as previously described (*4*, *14*, *15*, *29*). Briefly, 800 mL of *S. cerevisiae* containing one of the eleven Twin-StrepII-tagged importins were inoculated with a starting OD(600) of 0.035 in YPD. Cells were grown at 30 °C and 160 rpm to an OD(600) of 0.5-0.6. Afterward, cells were harvested by rapid filtration onto 0.45 µm nitrocellulose (Biorad), scraped off the membrane, and flash frozen in liquid nitrogen. Subsequently, cells were supplemented with 2.4 mL of lysis buffer (20 mM Hepes-KOH, pH 7.5, 140 mM KCl, 10 mM $MgCl_2$, 1 mM PMSF, 0.01 % IGEPAL, 0.1 mg/mL CHX, 1 tablet of cOMPLETE protease inhibitor per 50 mL) and lysed in a cryomill at 30 Hz for 2 min. Next, 2.4 mL of lysis buffer was placed into a 5 mL beaker on a magnetic stirrer and supplemented with 30 µL of 250 mM DSP. Gradually, the first half of the lysate powder was stirred in until thawed. Supplementation of DSP was repeated and the second half of lysate was added. Crosslinking was carried out for 10 min at room temperature while constantly stirring. The reaction was quenched by adding 400 µL of 2 M Tris-HCl pH 8.0. Next, the crosslinked lysate was cleared at 15,000 *g* for 3 min at 4 °C. The supernatant was transferred and absorbance at 260 nm of a 1:100 dilution was measured. To generate ribosome-protected footprints, 60 U [RNase I]/ absorbance [RNA] unit was added to each sample. RNase I digest was carried out by end-to-end mixing at 4 °C for 30 min. RNase I reaction was quenched by adding 200 U of Superase•In. The supernatant was then applied onto a 25 % sucrose cushion (25 % w/v sucrose, 20 mM Hepes-KOH, pH 7.5, 140 mM KCl, 10 mM $MgCl_2$, 0.01 % IGEPAL, 0.1 mg/mL CHX, 1 tablet of cOMPLETE protease inhibitor per 50 mL). Ribosomes were pelleted at 150,000 *g* for 2.5 hr. Then, the supernatant was discarded and the pellet was resuspended in 1 mL wash A buffer (20 mM Hepes-KOH, pH 7.5, 140 mM KCl, 10 mM $MgCl_2$, 0.01 % IGEPAL, 0.1 mg/mL CHX, 1 tablet of cOMPLETE protease inhibitor per 50 mL). 100 µg of RNA was collected after resuspension representing the "total translatome". This RNA was supplemented with 750 µL of 10 mM Tris HCl pH 8.0, and frozen in liquid nitrogen until further purification (see below).

The residual resuspended pellet was added to 250 µL of pre-equilibrated Streptactin resin (IBA). Additionally, 30 µL of BioLock (IBA) was added. Affinity purification of the Twin-StrepII tagged importin was carried out by end-to-end mixing for 1 hr at 4 °C. Next, beads were centrifuged at 500 *g* for 5 min and the supernatant was removed. Subsequently, beads were washed three times in 1 mL of wash A, each time applying a 1 min end-to-end mixing step. Last, beads were washed in wash B buffer (20 mM Hepes-KOH, pH 7.5, 140 mM KCl, 10 mM $MgCl_2$, 10 % v/v glycerol, 0.01 % IGEPAL, 0.1 mg/mL CHX, 1 tablet of cOMPLETE protease inhibitor per 50 mL) first time for 1 min and the second time for 4 min using end-to-end mixing.

After washing, the supernatant was removed and the beads were resuspended in 500 µL 10 mM Tris HCl pH 8.0 and 40 µL of 20 % SDS and gently mixed. Next, 750 µL of pre-warmed phenol-chloroform-isoamyl alcohol (PCI, 65 °C, Invitrogen) was added. This reaction was incubated at 65 °C, 1,400 rpm for 5 min and was immediately snap cooled on ice for 10 min prior to centrifugation at 15,000 *g* for 10 min. Once centrifuged, the aqueous phase was transferred into a new tube and supplemented with 750 µL of PCI. The mixture was occasionally vortexed at room temperature for 5 min and centrifuged again.

To remove residual PCI, a diethyl ether (Sigma Aldrich) wash was carried out. The residual organic solvent was evaporated using a Speedvac (Eppendorf).
RNA was precipitated by adding 3 M NaOAc, pH 5.5 to obtain a final concentration of 0.3 M, 2.5 μL of Glycoblue (Invitrogen), and equal volumes of isopropanol. Precipitations were vigorously vortexed and incubated at -80°C overnight. RNA was pelleted at 15,000 *g* and 4 °C for 90 min. the supernatant was removed and the pellet was washed three times with 70 % ethanol. The resulting RNA pellet was dried in the Speedvac (Eppendorf) and resuspended in 20 μL of 10 mM Tris HCl, pH 8.0.

**Library preparation**

Purified RNA and the 5' 6FAM labeled RNA marker were mixed with an equal volume of 2 x RNA loading dye (Thermo Scientific) and heat-denatured at 80°C for 2 min and immediately put back on ice before loading onto a pre-warmed 15 % denaturing urea polyacrylamide gel electrophoresis (PAGE) (Carl Roth). Gels were run for 3.5-4 hrs at 16 W until bromophenol blue emerged, disassembled, and stained in SybrGold (Invitrogen) according to the manufacturer's protocol and imaged using the Amersham Typhoon (GE Healthcare). RNAs at a size of 26-34 nt were isolated and crushed and soaked in 500 μL of Tris-HCl pH 8.0 at 70°C for 10 min and constant and vigorous shaking at 1,400 rpm. The slurry of gel pieces and buffer were then transferred into a Spin-X cellulose acetate column (0.22 μm; Corning), and centrifuged for 15 min and 4°C at 15,000 *g*. Flow through containing the RNA of interest was supplemented with 50 μL 3 M NaOAc, 2.5 μL Glycoblue co-precipitation agent (Invitrogen), and 500 μL isopropanol. The precipitation reaction was vortexed and incubated overnight at -80 °C. RNA was precipitated by centrifugation at 15,000 *g* for 90 min, washed three times with 70 % ethanol, and was finally resuspended in 15 μL nuclease-free water.
Following their purification, the ribosome footprints were dephosphorylated on their 5' and 3' end using 1 x FastAP buffer, 2 U FastAP (Thermo Scientific), and 20 U RiboLock (Invitrogen). Dephosphorylation was carried out for 15 min at 37 °C at 600 rpm. The reaction was quenched by heat-inactivation of the FastAP at 75°C for 5 min. Consecutively, 5' ends were phosphorylated by incubating the reaction with 20 U polynucleotide kinase (PNK; NEB), 1 mM ATP (Thermo Scientific), 1 x PNK buffer (NEB), and 20 U RiboLock to prevent RNA degradation. The phosphorylation was carried out at 37°C for 30 min.
Once the RNA was appropriately modified, integrity and concentration of the ribosome footprints were determined using the RNA Pico 6000 Assay Kit of the Bioanalyzer 21000 system (Agilent Technologies). 1 ng of RNA was used as input for library preparation using the NEXTflex Small RNA-seq Kit v3 (Perkin Elmer). Following this procedure, the quality of the libraries was assessed using the DNA High Sensitivity kit (Agilent Technologies), and the concentration of the library was determined using the Qubit DNA High Sensitivity kit using the Qubit 2.0 Fluorometer (Life Technologies). The concentration of each library was calculated and pooled at an equimolar amount. These multiplexed pools were finally purified with SPRI select beads with an excess of 1.3 x of beads (Beckman Coulter). The purified ribosome profiling library pool was loaded onto the Illumina sequencer NextSeq2000 and sequenced uni-directionally, yielding ~ 1,379 million reads with a size of 72 bases.

**Sequence processing**

Data from the NextSe q2000 was processed following instructions published in Galmozzi et al. (*14*) and the script suite provided within the aforementioned instructions (https://doi.org/10.5281/zenodo.2602493). Reads were cleaned and trimmed using cutadapt (v2.3) (*65*). Non-coding RNAs of *Saccharomyces cerevisiae* were filtered and excluded for further analysis using a non-coding reference genome (R64-1-1.ncrna). Reads encoding for coding RNAs were mapped to a *Saccharomyces cerevisiae* (R64-1-1) reference genome using tophat2 (v2.0.10). Ribosome centering was carried out as described within the script suite (https://doi.org/10.5281/zenodo.2602493).
Output files of script A which contain the numbers of reads per genomic position were extracted and used as input for our MATLAB scripts (v9.7.0.1296695 (R2019b) Update 4) as described in Seidel et al. (*29*) and for our novel python pipelines.

### Ribosome profiling data analysis and target identification

Genes were mapped to the S288C (R64) reference as downloaded from SGD (*66*) (*S288C_reference_sequence_R64-1-1_20110203.fsa*); introns and exons were mapped to sequence data using *sacCer3.ensgene.gtf* as extracted from Ensembl (*67*). The following analysis is based on normalized read counts as the output of the aforementioned sequence processing. Per experiment (IP, and total, alike) replicates were averaged for each nucleotide. In each experiment, a mean correlation value (Pearson) was calculated for each gene across replicates; this is later used for quality filtering of potential pulldown targets (threshold 0.6). Gene ribosome profiles for each replicate, and averaged, were smoothed using a sliding window of 100 nucleotides (calculation of moving average) using the *pandas* (Python) package (*68*). The orientation of the respective strand was considered; introns were not removed for smoothing. From these smoothed profiles in pulldown experiments (IP) and respective controls (total), the area under the curve (AUC) was calculated (excluding introns) using the trapezoidal rule as implemented in the *numpy* (Python) package (*69*). Coverage levels of each gene (in percentage) are calculated from smoothed profiles, for both pulldowns and controls.

For further target identification, transposon-related genes were removed from quantification to avoid bias in the analysis. The fold change between pulldown and control (IP/total) was calculated for each gene profile by calculating the ratio of respective vectors; the fold change of AUC-values was also collected for each gene. These fold change gene profiles will further be used in the target identification, and are illustrated in figures (see **Figs. 1**, **2,** and **5**; **Figs. S7, S9, S10**) and labeled as “Enrichment (IP/total)”. We approached target identification using an FDR calculation based on a true positive set of targets as has been defined for Srp1 (see list in **Table S1**) from the literature. Prior to FDR calculation, the following filtering steps were applied for each gene profile: i) gene coverage is to be set at a minimum of 80 %, ii) the AUC-value in the pulldown condition has to be larger than 5 (i.e. removing signal from lower quantile, ), and iii) the AUC derived from the fold change profile in the pulldown condition has to at least correspond to the AUC derived from the respective fold change profile in the no-bait wildtype condition (**Fig. S3**). The latter condition puts profiles from pulldown experiments in relation to the no-bait wildtype conditions for the first time in the analysis pipeline. For the FDR calculation, the AUC-value per gene and pulldown was scaled relative to the no-bait wildtype condition, i.e. for each gene the AUC-values are summarized for the pulldown and no-bait wildtype experiment; the gene-specific AUC-value in the pulldown is then divided by the latter sum and reflected as a percentage (e.g. shown in **Fig. 2A**). This no-bait wildtype-scaled AUC-value was then used for calculating a cut-off threshold at FDR 1% (based on the true positive set as mentioned above). At a cutoff-threshold of 72%, most of the true positive targets of Srp1 can be recovered (see ROC Curve, **Fig. S3B**). The resulting target set for other pulldowns was then further subjected to manual inspection, and a mean correlation threshold of 0.6 (across replicates) was set for each gene for increasing the quality of hit profiles.

We additionally conducted an analysis using the DESeq2 R-package (*70*) on the smoothed normalized counts of the genome profiles across pulldowns relative to the no-bait (wildtype) control. For each pulldown analysis, 4 replicates from the selective translatome were considered, as well as the 4 replicates from the no-bait control (factored as conditions). The resulting log2 fold changes and adjusted *P*-values (following the default Benjamini-Hochberg adjustment) are illustrated in **Fig. S5**.

To map SeRP profiles, we extended our previously published scripts (*29*). The updated script suit will be made available on Zenodo.

### Protein analysis

To analyze the bait-StrepII enrichment and the efficiency of crosslinking, samples were mixed with 4 x NuPAGE loading dye containing 5 % of beta-mercaptoethanol if not otherwise specified. Samples were boiled at 70 °C for 5 min prior to loading. Loading of the NuPAGE Bis-Tris gels (MW < 100 kDa) (Invitrogen) was conducted as follows: four microliters of the lysate (~0.08 %) for reducing and non-reducing conditions (non-reducing: 4 x NuPAGE loading dye without beta-mercaptoethanol) were loaded to address crosslinking efficiency; four microliters of the resuspended ribosome pellet (0.4%) and 10 µL of the boiled Streptactin resin (equivalent to 4.0 %) were loaded to evaluate bait-StrepII enrichment. The samples were run in MOPS buffer (Invitrogen) at 180 V for 50 min.

To analyze the crosslinking efficiency, gels were stained with Instant Blue (abcam) as stated in the manufacturer’s protocol.

For the analysis of the bait-enrichment, the protein was transferred onto 0.45 µm TransBlot Turbo nitrocellulose (Bio-Rad) using the manufacturer's High MW setting of the TurboBlot (Bio-Rad). Subsequently, membranes were blocked in 5 % milk in TBS-T (0.02% Tween-20) for 1 hr at room temperature while shaking. Afterward, the membranes were incubated with the monoclonal anti-StrepII antibody (EPR12666; ab180957; Lot. No. GR3212622-7) diluted in 1:5,000 in 5 % milk in TBS-T. The membranes were incubated with the primary antibody at 4 °C overnight. Afterward, membranes were vigorously washed three times with TBS-T prior to applying the secondary antibody. The secondary anti-rabbit IgG, HRP-linked antibody (Jackson ImmunoResearch) diluted 1:10,000 in TBS-T was incubated for 1 hr at room temperature while shaking. The previously described washing procedure was repeated and the protein was detected by applying Clarity Max Western ECL solution (Bio-Rad) and imaged using the Chemidoc (Bio-Rad).

**Confocal imaging**

Cells were grown overnight in a synthetic complete medium without uracil (SC -Ura; ForMedium) at 30 °C at 180 rpm. Before imaging, cells were diluted to OD(600) 0.2 and again grown under the aforementioned conditions until OD(600) 0.4-0.6. 100 µL of the cell suspension was placed onto Concanavalin A (1 mg/mL, Sigma Aldrich) coated slides, settled for 5 min, and washed three times in SC -Ura.

Live-imaging was performed on a laser scanning confocal microscope Leica Stellaris 5. Cells were imaged at room temperature. Images were acquired using a 63 X/ 1.2 NA water objective (HC PL APO CS2). Z-stacks were collected using the following image setup: excitation (GFP): 488 nm; emission: 493-591 nm; zoom factor 2, four line averaging, and in bi-directional mode. The microscope was operated using the LAS X software provided by Leica Microsystems CMS GmbH (v.4.4.0.24861). 9-11 images per z-stack were maximum intensity projected using ImageJ (v.1.52).

**GO enrichment analysis**

GO-categories from Cellular Compartment (cc), Molecular Function (mf), and Biological Processes (bp) were extracted from Uniprot (https://www.uniprot.org/), along with mappings of yeast proteins to GO-categories. For the analysis, only GO-terms with more than 100 protein entities were considered. The Fisher enrichment calculation (using the *scipy* Python package) was applied to calculate the odds enrichment of each GO-term within the specific hit set relative to all quantified proteins in the Ribosome Profiling dataset. For visualization, only GO-terms were considered that were significant ($P \leq 0.1$) in at least one pulldown condition. *P*-values were not adjusted. For visualization, the odds-matrices were hierarchically clustered using the Euclidean metric (*seaborn* Python package).

**Analysis of pI-values**

Uniprot IDs of the targets and nuclear proteome (GO: 0005634) were fetched on Expasy (https://web.expasy.org/compute_pi/). pI-values were extracted and displayed as a violin plot using GraphPad Prism (v9.0.0). Non-parametric (two-sided) Mann-Whitney *U*-test has been applied to calculate the difference in the pI distributions across conditions. *P*-values were adjusted using Benjamini-Hochberg.

**Analysis of the domain-onset relationship**

Domain architecture of respective proteins was analyzed using InterPro (https://www.ebi.ac.uk/interpro/) (*71*) and Pfam domain descriptions (*72*). Additionally, disorder predictions generated by InterPro were considered. Note that Nup1 and Nup60 were annotated based on their recent domain diagrams presented in Mészáros et al. (*73*). "Domain boundaries" reflect proteins with 50 amino acids off from the domain end or start. If domains are known for several functions, the domain was assigned to both of its functions. Functions of the respective domains were assigned manually according to the Pfam entry and availability of structures embedded in Pfam.

**Meta-analysis regarding Figs. S10 and S11**

Yeast proteins were mapped to data derived from several publications, including Ghaemmaghami et al., 2003 (protein abundance) (*74*), Arava et al., 2003 (protein synthesis rates) (*75*), and Wilmund et al., 2013 (Ssb1/2-dependencies) (*45*). For calculations regarding protein synthesis and protein abundance, mapped data was stratified across pulldown conditions. A Kolmogorov-Smirnov test (two-sided) was applied for comparing specific pulldown conditions with the data assigned to all nuclear proteins. *P*-values were adjusted using Benjamini-Hochberg. For Ssb2 and essentiality mapping, we applied Fisher enrichment calculation- again relative to the nuclear proteome.

**Prediction of NLS motifs using AlphaFold**

We searched for putative NLS motifs in the target proteins by running AlphaFold-Multimer (*35*) structure prediction for Srp1 (AA 70-542, without the autoinhibitory N-terminal peptide) in a complex with consecutive 100 AA. long fragments spanning the target sequences from the N-terminus to residue 550 with 50 AA. overlaps. AlphaFold-Multimer was run with default parameters except the max_recycles parameter set to 12 (to ensure convergence of the modeling). The predictions were scored according to combined ipTM (interface predicted TM-score) and pTM score (predicted TM-score), as returned by AlphaFold, and the top-scoring model for each pair was taken for further analysis (**Table S3**). To calculate the multiple sequence alignment in **Fig. 3B**, we selected all predictions that i) exhibited the ipTM+pTM score > 0.7, ii) bound to the canonical NLS-binding site in Srp1, iii) bound in the N- to C-terminus orientation as known from crystal structures of Srp1-NLS complexes. The selected structures were superposed using UCSF Chimera (*76*) and the multiple sequence alignment was derived from the superposition. Figures were prepared using UCSF ChimeraX (*77*) and Jalview (*78*).

**Multiple sequence alignment**

Protein sequences for the 11 importins were retrieved on UniProt. FASTA-files were used for multiple sequence alignments Clustal Omega (https://www.ebi.ac.uk/Tools/msa/clustalo/). The dendrogram was generated upon the multiple sequence alignment output in Clustal Omega.

**Quantification and statistical analysis**

Significance levels are shown when $*P< 0.05$, $**P< 0.01$, $***P< 0.005$, $****P< 0.001$. For adjustments, we applied the Benjamini-Hochberg method as implemented in the Python *scipy* package. We considered *P*-values as not significant (ns) when $P> 0.05$. All statistical tests were applied in a two-sided manner (if not indicated otherwise) and consider the underlying value distributions. Accordingly, parametric or non-parametric tests have been applied.

Selective ribosome enrichment plots depict data of $n$=4 replicates for each SeRP experiment. Arbitrary background thresholds indicated as a grey dashed line was set to 1.5 or 2.0, respectively, and are specified in the respective figure legends.

Western Blot analysis to analyze the enrichment of importins and Coomassie gels to check for crosslinking experiments shown in **Figs. S1C** and **S1D** were performed for every biologically independent experiment ($n$=4).

Imaging shown in **Figs. 2G** and **2H** were repeated at least twice.

**Data and code availability**

Underlying metadata including uncropped images of Western Blots and Coomassie gels, data used of previously published data (metadata), and data of Ssb1/2 (*17*) and TRiC (*16*) will be made available in a **Source Data file** upon publication.

The selective ribosome profiling data for all experiments conducted in this study are available on the European Nucleotide Archive database under the accession code PRJEB53855 (accessible for reviewers upon request)**.** For the initial processing of the ribosome-protected footprints, we used a script suite for selective ribosome profiling (*14*) which can be accessed on Zenodo [https://doi.org/10.5281/zenodo.2602493]. This script suite also provides the required reference genome files for *Saccharomyces cerevisiae* (*S288C_reference_sequence_R64-1-1_20110203.fsa*), as downloaded from SGD [http://sgd-archive.yeastgenome.org/sequence/S288C_reference/genome_releases/S288C_reference_genome_R64-1-1_20110203.tgz]. We also used sacCer3.ensgene.gtf for mapping introns and exons, as extracted from Ensembl (*67*) [https://hgdownload.soe.ucsc.edu/goldenPath/sacCer3/bigZips/genes/sacCer3.ensGene.gtf.gz].

Previously published SeRP data for Ssb1/2 (*17*) and Ssb and TRiC (*16*) can be accessed at Gene Expression Omnibus (GEO) under GEO: GSE93830 [https://www.ncbi.nlm.nih.gov/geo/query/acc.cgi?acc=GSE93830] and GEO: GSE114882 [https://www.ncbi.nlm.nih.gov/geo/query/acc.cgi?acc=GSE114882].

The structure of the autoinhibitory NLS of Srp1 bound to Srp1 (**Fig. 3A**) can be found in the Protein Data Bank under the accession number PDB: 1WA5 [10.2210/pdb1WA5/pdb] (*37*). AlphaFold-Multimer models will be made available upon publication and can be requested by the reviewers. AlphaFold models of the r-proteins (**Fig. 5**) originate from the AlphaFold database (https://alphafold.ebi.ac.uk/) (*46*, *47*).

Our previously published MatLab scripts are deposited in Zenodo (DOI: 10.5281/zenodo.5887402). The updated version and the SeRP hit identification pipeline and the AlphaFold analysis pipeline will be made available on Zenodo upon publication and can be requested by the reviewers in advance.

### Supplementary Figures

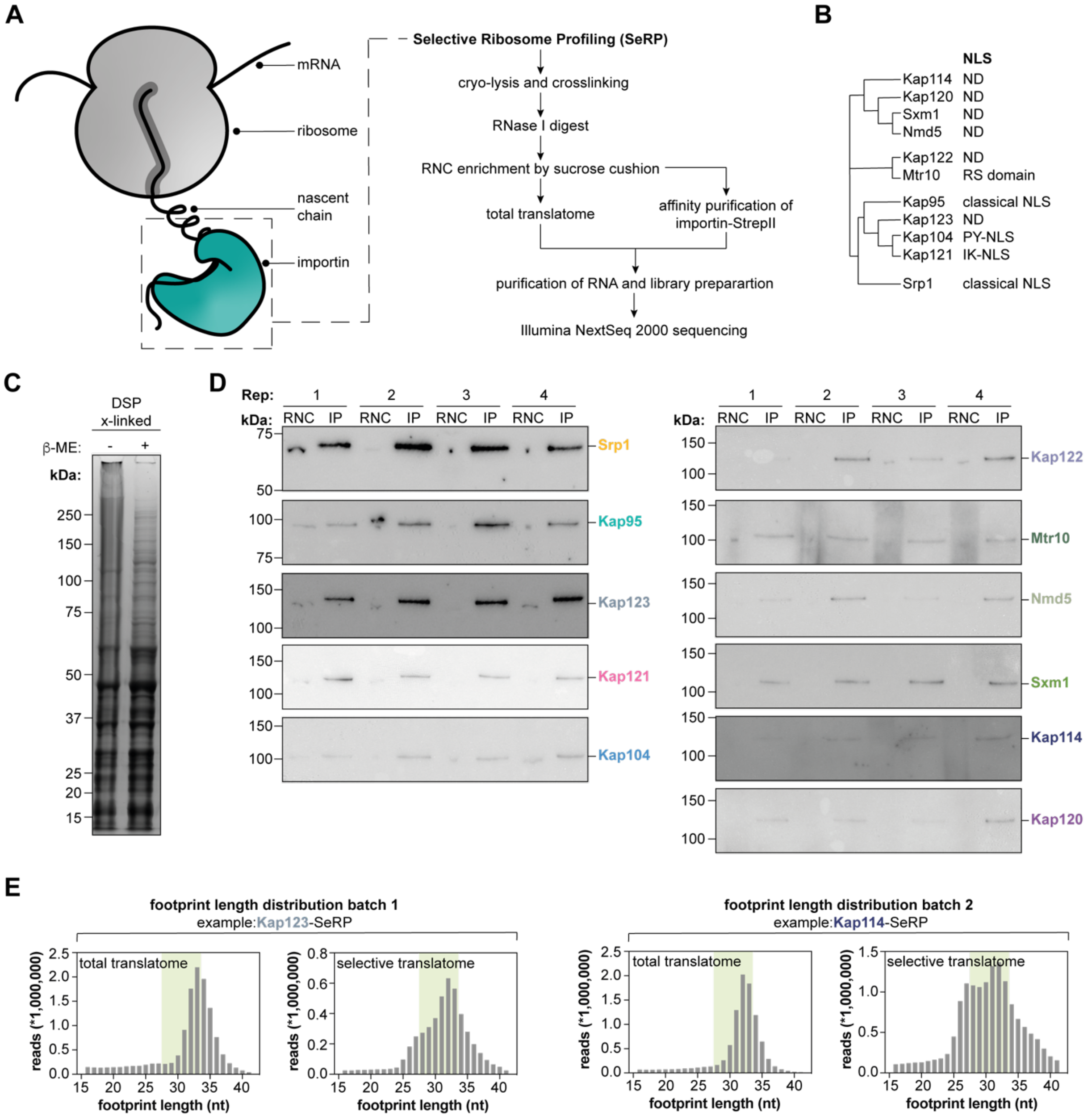

**Figure S1: A,** Enrichment strategy for importin-bound nascent chains and selective ribosome profiling procedure. **B,** Multiple sequence alignment of importins depicts three distinct groups of importins, consensus sequences in *S. cerevisiae* are indicated if applicable. **C,** Analysis of cross-linking efficiency. To stabilize interactions, lysates were cross-linked in presence of DSP. Cross-linked lysate shows an upwards shift of protein bands and results in a smear. Reduction of DSP using reducing agents such as β-mercaptoethanol (β-ME) validates that cross-linking was successful under the relevant conditions. Protein was visualized by Coomassie Blue. **D,** Western blot analysis of the individual pulldown experiments. StrepII-tagged importins could be enriched from the ribosome-nascent chain (RNC) fraction. StrepII-tagged protein was detected using an anti-StrepII-tag antibody. **E,** Footprint length distribution of the total translatome and the selective translatome channel. Distribution shows average read distribution across $n$=4 replicates. ND: not determined; NLS: nuclear localization sequence; x-linked: crosslinked; β-ME: β-mercaptoethanol; Rep: replicate; IP: immunoprecipitation.

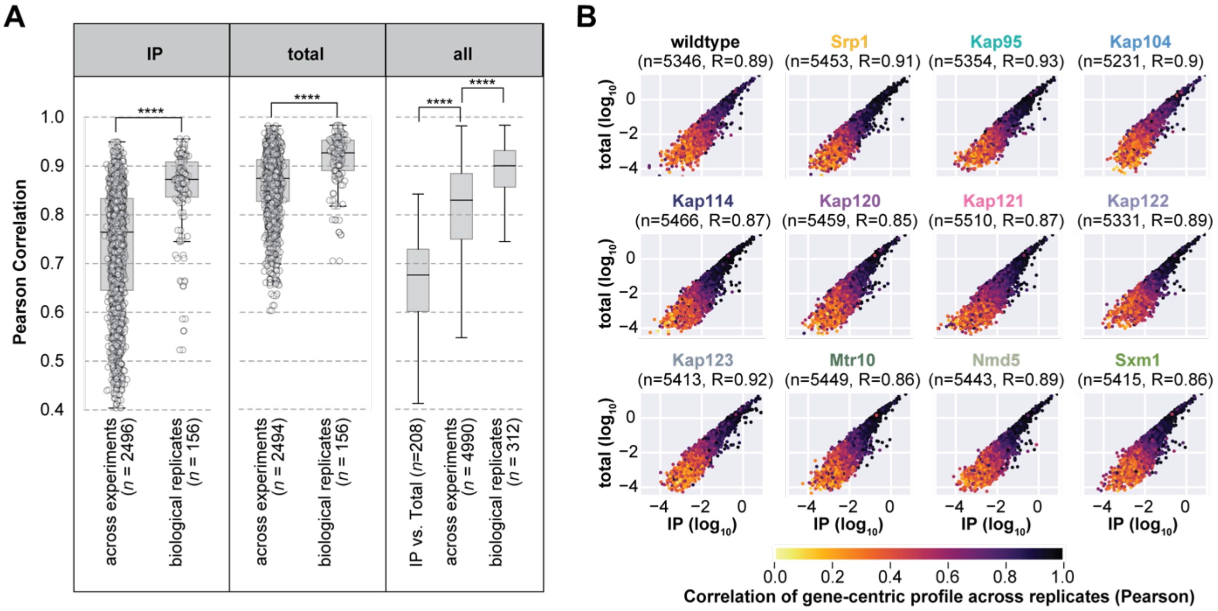

**Figure S2: A,** Box plots indicating the distribution of the Pearson correlation coefficients as calculated from normalized counts across experiments and replicates. The central line in the box plots indicates the median, the bottom and top edges of the box the IQR, and the box plot whiskers represent 1.5 times the IQR. For comparing medians of distributions, a two-sided t-test was applied, and significance is indicated (all P-values < 0.001,****). **B,** Scatterplot showing averaged normalized count in IP (x-axis) and total translatome (y-axis) in log10-scale. Each dot represents a gene; the color indicates the Pearson correlation of the gene-centric profile across replicates (Pearson).

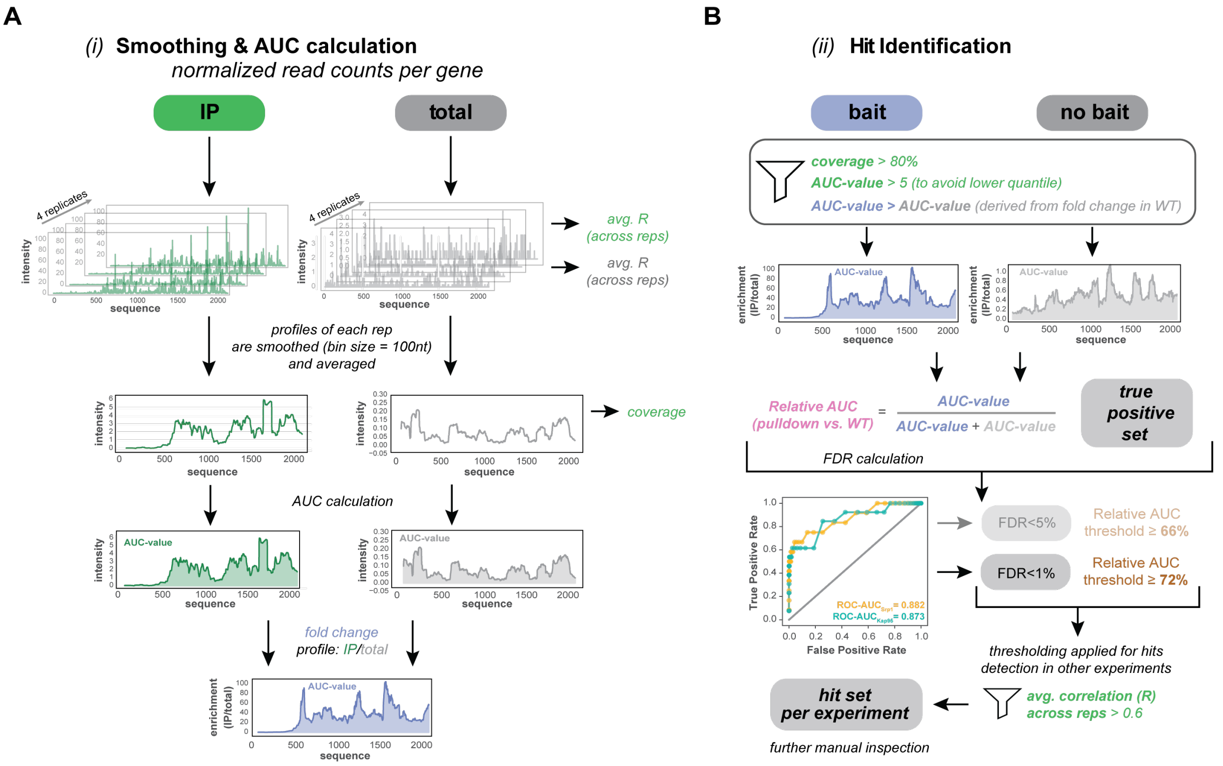

**Figure S3:** Hit detection pipeline developed for this study. **A,** Calculation of individual selective ribosome profiles. IP and total translatome (total) were averaged across the four replicates and smoothed with a bin size of 100 nt. Resulting profiles were used to determine the coverage and the area under the curve (AUC)-value in the IP and total, respectively. To obtain a fold change profile or enrichment profile (blue), the IP profile (green) was divided by the associated total profile (grey). **B,** Hit identification across all experiments. The obtained enrichment profiles were filtered according to coverage and AUC-value (see **Materials and Methods** for a more detailed explanation). Each gene-centric enrichment profile is put into context with its corresponding profile in the wildtype condition (no bait), hence a wildtype-relative AUC can be calculated. This relative AUC-value is used for false discovery (FDR) calculation based on the true positive set as defined in **Table S1**. A Receiver Operating Curve (ROC) is calculated, indicating the recovery of true hits within the data. At a threshold of 72% (Relative AUC), an FDR of <1% is achieved and hence can be used for filtering highly reliable hits. The final hit set per experiment can be further filtered and manually curated.

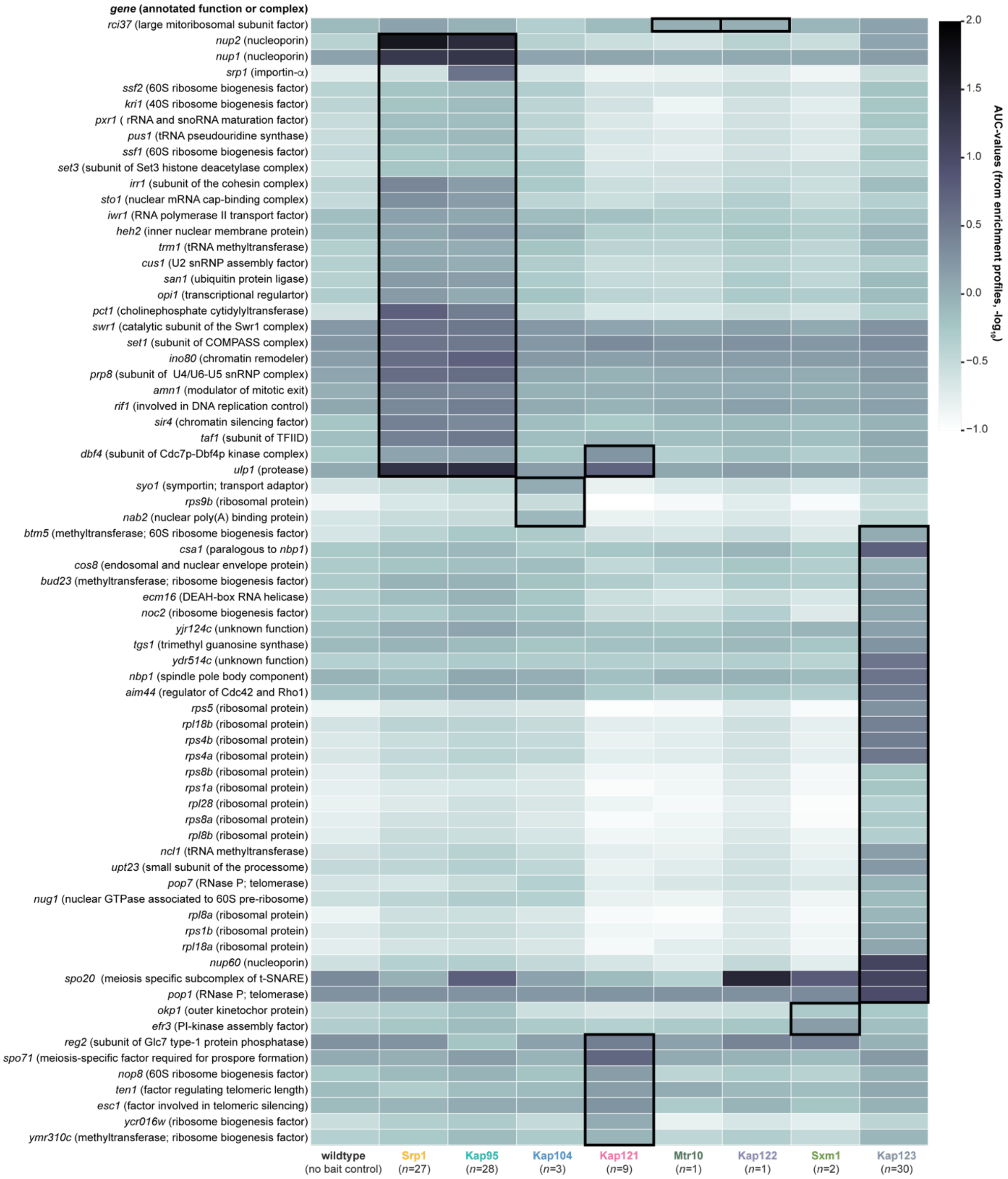

**Figure S4:** AUC-heatmap of all reliably identified hits across different experiments. AUC-values are presented and can be compared to the AUC-values generated in the wildtype (no bait control). Circulated genes represent hits within their respective SeRP experiment.

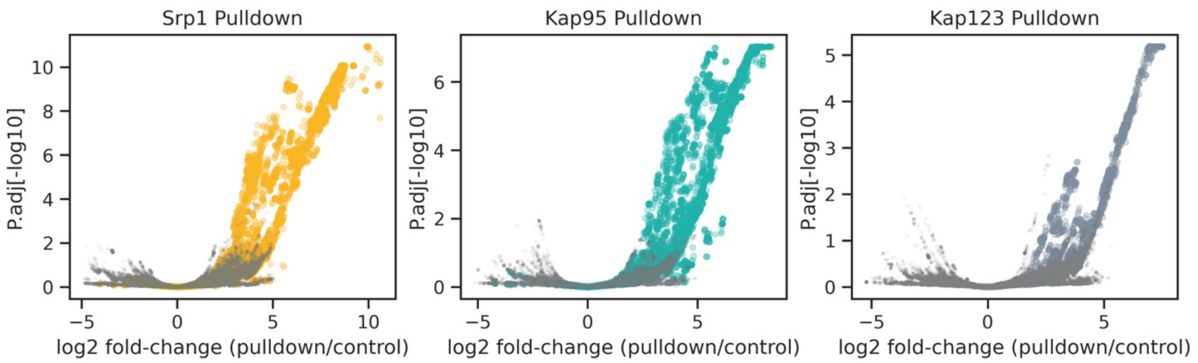

**Figure S5:** Volcano plots illustrating DESeq2-derived results for profiles (normalized counts) across the entire genome for respective pulldown relative to wildtype (no-bait) control (see corresponding **Materials and Methods** section). The x-axis indicates the log2 fold change, and the y-axis the adjusted *P*-value in a –log10 scale. Each dot corresponds to a nucleotide position (hence strong cross-correlation in analysis); grey dots signify positions within genes that have not been identified as targets of the respective importin our pipeline, whereas colored dots correspond to genome positions that reside within target or cargo genes. The DESeq2 analysis demonstrates that normalized counts in the identified cargo genes are significantly enriched relative to the no-bait control (Fisher exact test shows $P < 0.0001$ for all three pulldowns). This Supplementary plot shows the underlying data for Srp1, Kap95, and Kap123, where we also found the most reliable hits. The other pulldowns did not demonstrate a significant signal in the DESeq2 analysis.

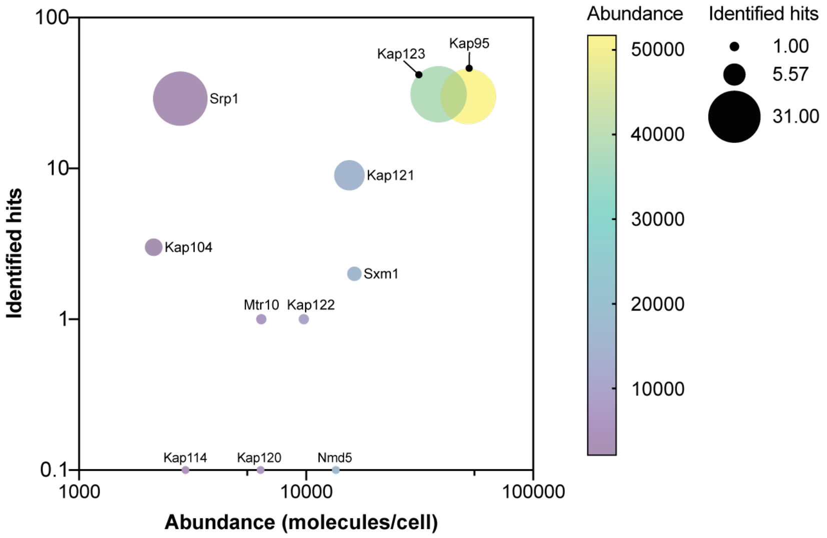

Source: Ghaemmaghami et al, 2003, *Nature.*

**Figure S6:** Relationship of importin abundance and number of identified targets. The analysis reveals that the number of identified hits does not depend on the abundance of the respective importin. Bubble size represents the number of identified hits. Abundance of the respective importin is encoded by a color gradient. Abundances of importins (molecules/cell) were extracted from Ghaemmaghami et al. (*74*).

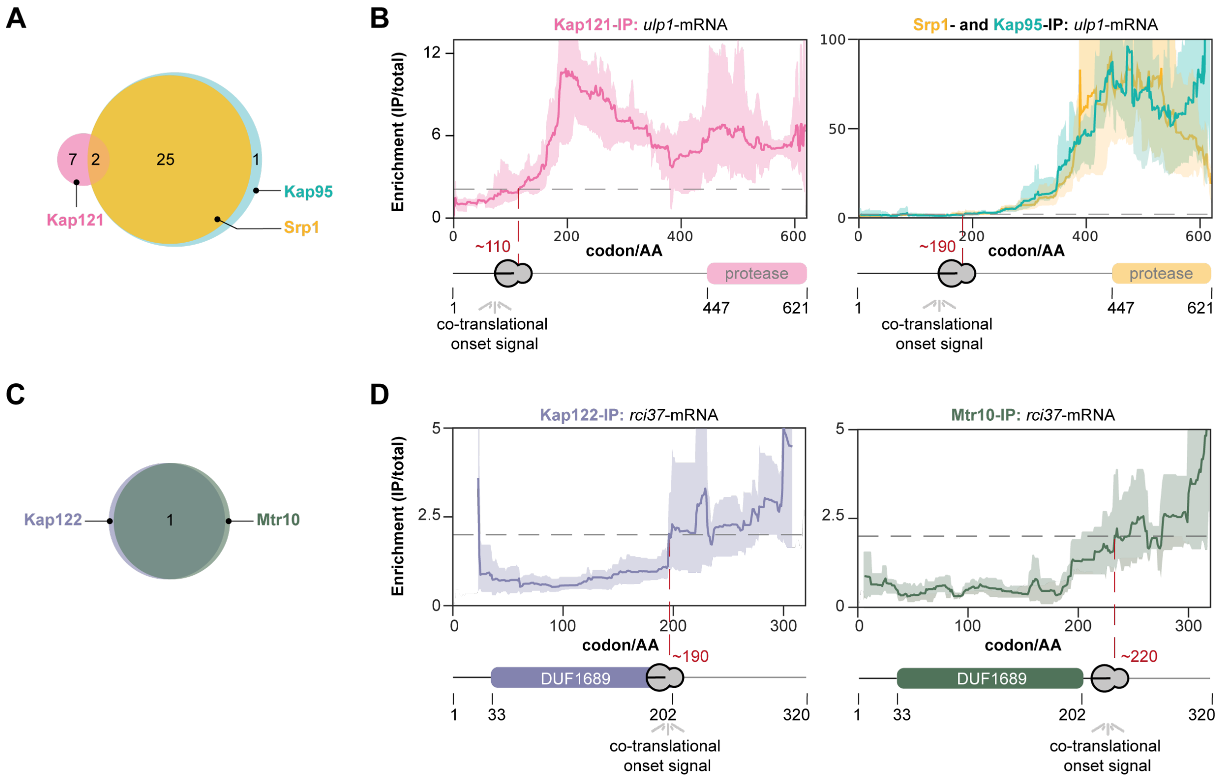

**Figure S7:** Co-translational interactions show minor redundancy. **A,** and **B,** Venn-diagram indicating overlap of the co-translational cargoes between Kap121 and Srp1/Kap95. They share Ulp1 and Dbf4 as substrates but the apparent onset is different implying that Kap121 pioneers in binding to nascent Ulp1. **C,** and **D,** Kap122 and Mtr10 both capture the nascent chain of Rci37. SeRP profiles (IP/total) are shown for the respective mRNA targets from n=4 biologically independent replicates (solid lines are averaged across replicates; shades reflect largest to smalls replicate value interval). Grey dashed lines indicate an arbitrary threshold of 2 used for onset estimation (red dashed line). IP: immunoprecipitation; AA: amino acid; DUF: domain of unknown function.

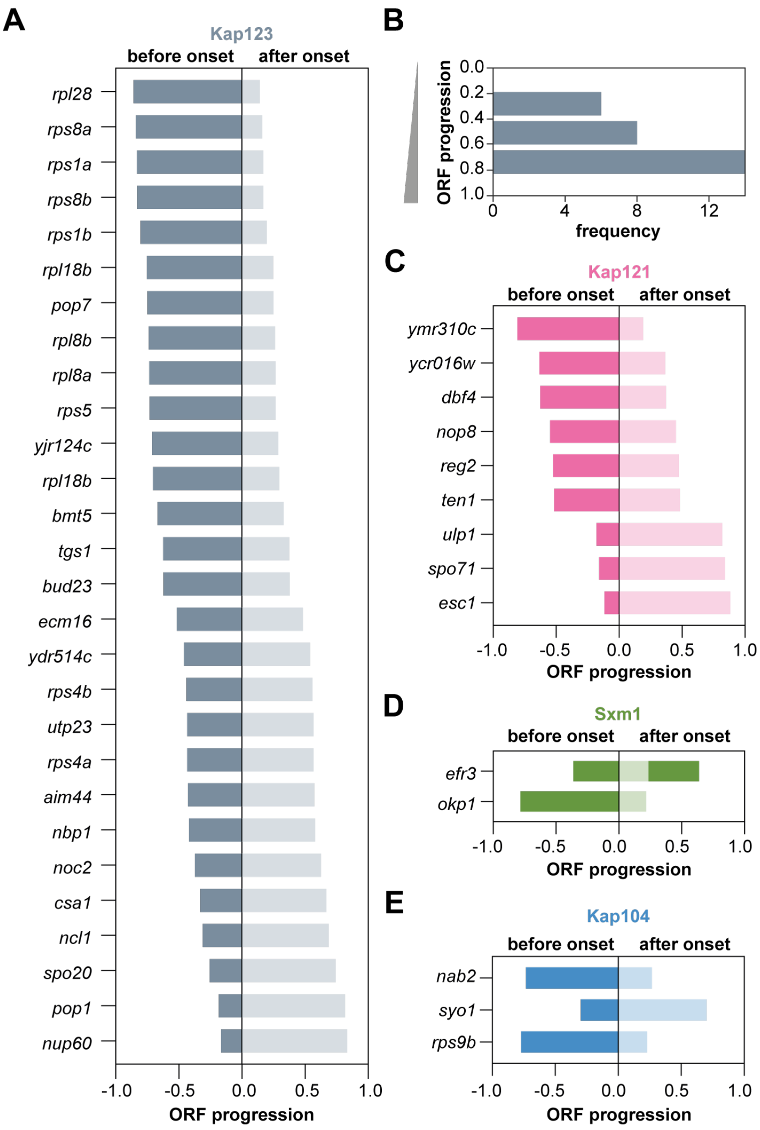

**Figure S8:** Onset distribution across importin-SeRP experiments other than Kap95/Srp1. **A,** Onset distribution for Kap123 cargoes. **B,** The respective histogram indicates a C-terminal bias. Onset plots for **C,** Kap121, **D,** Sxm1, **E,** Kap104. All onset plots show normalized ORF length for all genes. Apparent onsets are centered to zero. ORF: open reading frame.

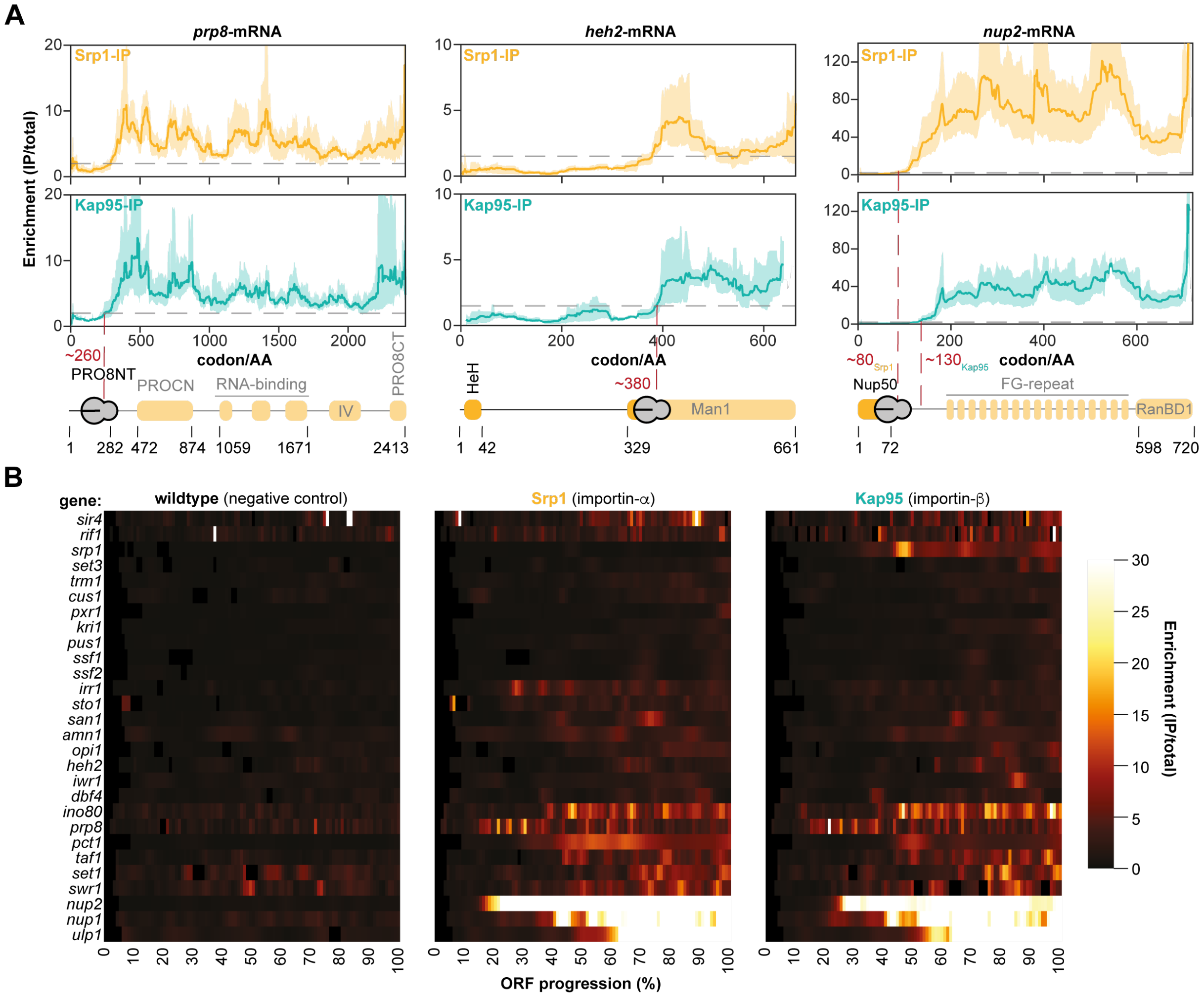

**Figure S9:** Synchronous binding of Srp1 and Kap95 on the nascent chains. **A,** Representative SeRP curves show that Srp1 and Kap95 mutually act on the nascent chains of their cargo in the case of nascent Prp8 and Heh2. In contrast, nascent Nup2 appears to be bound first by Srp1 and subsequently by Kap95. We note that Heh2 expresses an NLS between residues 102-137 (*79*) and requires an additional linker for nuclear localization (*80*, *81*). SeRP of Kap95 and Srp1 suggests binding of importin only upon exposure of the linker from the exit tunnel. SeRP profiles (IP/total) are shown for the respective mRNA targets from n=4 biologically independent replicates (solid lines are averaged across replicates; shades reflect largest to smalls replicate value interval). Grey dashed lines indicate an arbitrary threshold of 2 used for onset estimation (red dashed line). **B,** Heatmap representation of all Srp1 and Kap95 co-translationally targeted cargoes in relation to wildtype (no bait) negative control, showing that signals arise at similar regions within the respective ORFs, while almost no signal is generated in the no-bait control. Heatmap colors indicate the enrichment (IP/total) value according to the scale bar. Each profile and heatmap value were derived from n=4 biologically independent samples for each bait. IP: immunoprecipitation; AA: amino acid; HeH: helix-extension-helix domain; Man1: Man1-Src1p-C-terminal domain; FG-repeat: phenylalanine-glycine repeat; RanBD1: Ran binding domain 1.

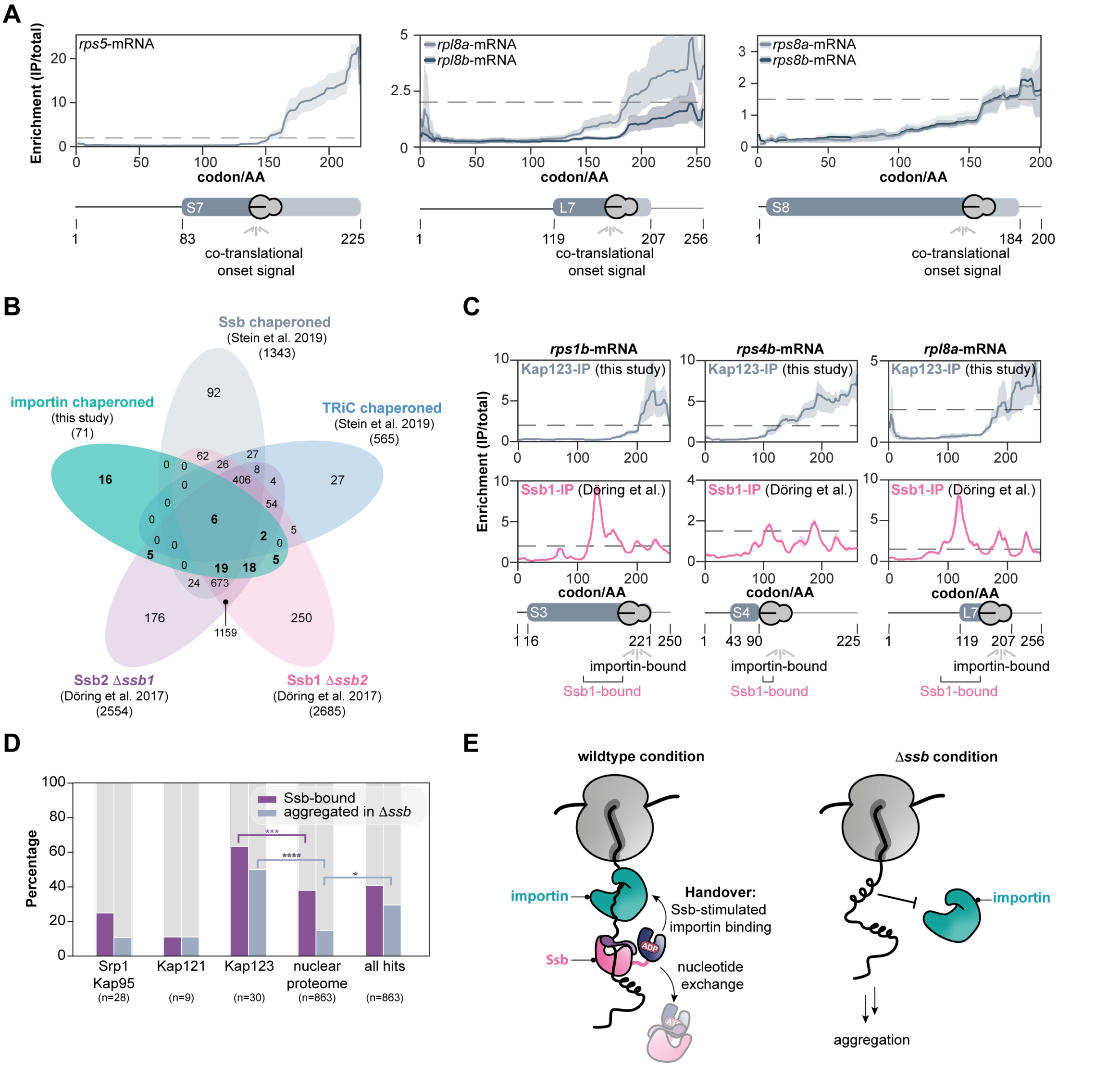

**Figure S10:** Ribosomal proteins captured by Kap123. **A,** Representative SeRP curves for ribosomal proteins. Curves for paralogues ribosomal proteins show similar shapes and onsets but their signal fluctuates according to their expression level. *rpl8a* expression is higher with respect to *rpl8b*, resulting in a stronger signal, while *rps8a* and *rps8b* are expressed at similar levels resulting in two almost identical curves. **B,** Venn-diagram of substrate overlap across Ssb-, TRiC- and importin chaperoned cargo. **C,** Extended evidence that Ssb1 binding to nascent chains precedes Kap123. SeRP profiles (IP/total) are shown for the respective mRNA targets from n=4 biologically independent replicates (this study), Ssb1-SeRP experiments originating from Döring et al. (*17*) represent profiles generated form n=2 biologically independent replicates (solid lines are averaged across replicates; shades reflect largest to smalls replicate value interval). Grey dashed lines indicate an arbitrary threshold of 1.5 or 2 used for onset estimation (red dashed line). **D,** Kap123 substrates are enriched for proteins that aggregate in the absence of Ssb2. ***$P$= 0.007 (Kap123 [Ssb-bound], nuclear proteome [Ssb-bound]); ****$P$< 0.001 (Kap123 [aggregated in Δ*ssb*], nuclear proteome [aggregated in Δ*ssb*]), *P= 0.036 (all hits [aggregated in Δ*ssb*], nuclear proteome [aggregated in Δ*ssb*]). ns $P$> 0.05, *$P$< 0.05, **$P$< 0.01, ***$P$< 0.005, ****$P$< 0.001. Fisher Exact test. **E,** Possible interpretation: under wildtype conditions, Ssb may maintain importin recognition sites degenerated allowing for importin binding. Within *ssb*Δ, nascent chains undergo folding thus masking the linear NLS and ultimately resulting in aggregation. IP: immunoprecipitation; AA: amino acid.

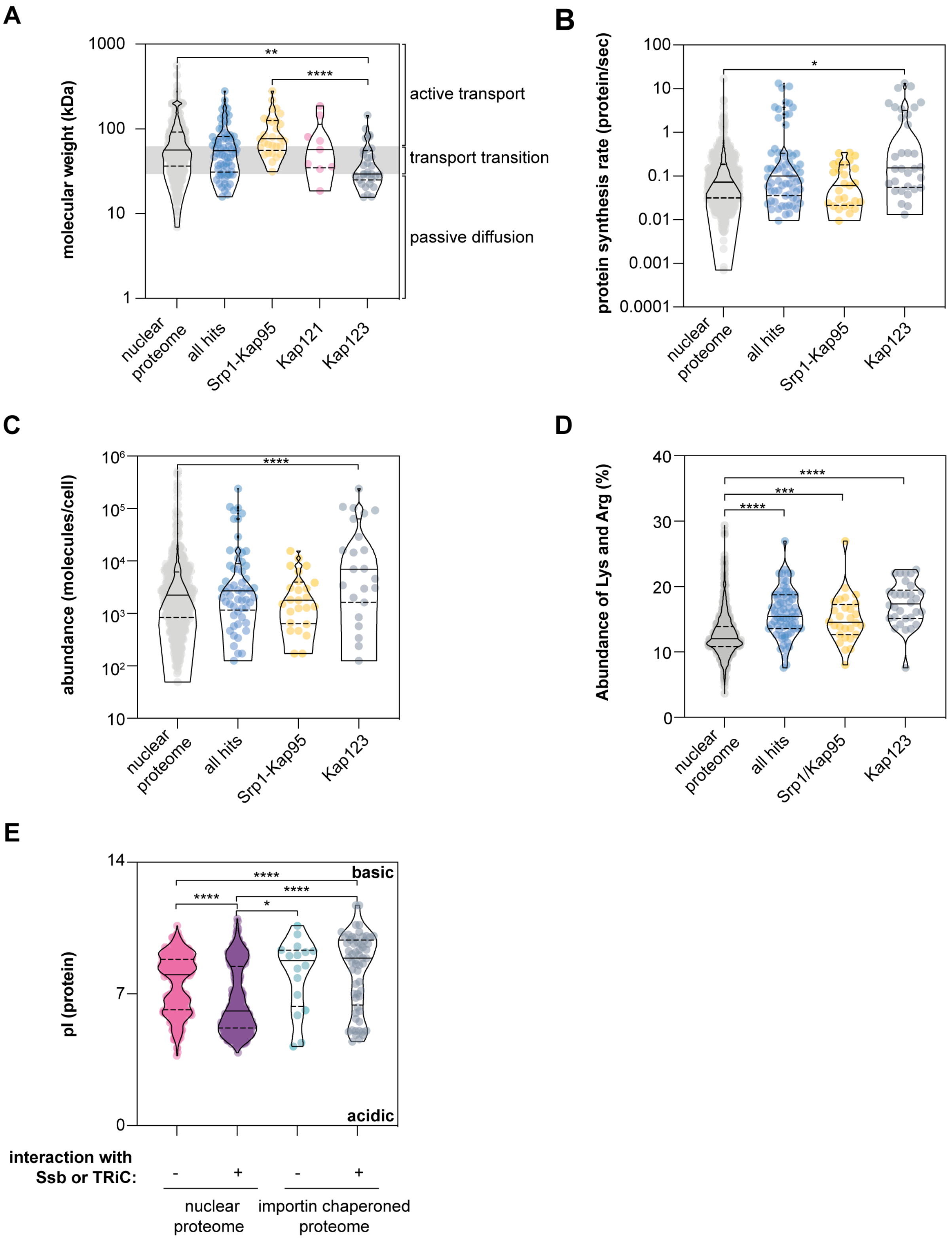

*Figure legend on the next page.*

**Figure S11: A,** Co-translational chaperoning of cargo is independent from cargo molecular weigth. Srp1-Kap95 and Kap121 chaperone cargo that tends to require active transport, while the median size of Kap123 cargo is below the size threshold of permeability barrier for passive diffusion. **$P$= 0.0085 (nuclear proteome, Kap123); ***$P$< 0.001 (Srp1-Kap95, Kap123). **B,** Kap123 cargoes are proteins with high protein synthesis rate. *$P$= 0.0227 (nuclear proteome, Kap123). **C,** Kap123 cargoes are highly abundant. ****$P$< 0.001 (nuclear proteome; Kap123). **D,** Proteins bound by importins show higher content of lysine (Lys) and arginine (Arg). "all hits" summarizes all hits across pulldowns. ****$P$< 0.001 (nuclear proteome, all hits); ***$P$= 0.0013 (nuclear proteome, Srp1-Kap95); ****$P$< 0.001 (nuclear proteome, Kap123). A two-sided Mann-Whitney U-test has been applied. **E,** Ssb and TRiC chaperones do not protect proteins with a high pI-value. ****$P$< 0.001 (nuclear proteom [non-chaperoned], nuclear proteome [Ssb-, TRiC-chaperoned]); *$P$= 0.0308 (nuclear proteome [Ssb-, TRiC-chaperoned]), importin chaperoned proteome [non-chaperoned]); ****$P$< 0.001 (0308 (nuclear proteome [Ssb-, TRiC-chaperoned]), importin chaperoned proteome [Ssb-, TRiC-chaperoned]); ***$P$< 0.001 (nuclear proteome [non-chaperoned], importin chaperoned proteome [Ssb-, TRiC-chaperoned]). "all hits" summarizes all hits across pulldowns. ns $P$> 0.05, *$P$< 0.05, **$P$< 0.01, ***$P$< 0.005, ****$P$< 0.001. Violin plots show median and quartiles. For all figures, the Mann-Whitney U-test has been applied to calculate corresponding $P$-values and account for variation in distributions. pI: isoelectric point.

### Supplementary Tables

| Cargo | Importin | Reference | Co-translational interaction detected |
|---|---|---|---|
| Nup1 | Kap95, Srp1 | shown in (*73*) | yes |
| Nup2 | Kap95, Srp1 | shown in (*73*, *82*) | yes |
| Ulp1 | Kap95, Srp1 | shown in (*57*) | yes |
| Asr1 | Kap95 | shown in (*83*) | no |
| Pct1 | Kap95, Srp1 | shown in (*84*) | yes |
| Heh2 | Kap95, Srp1 | shown in (*81*) | yes |
| Cdc45 | Kap95, Srp1 | shown in (*36*) | no |
| Vps75 | Kap95, Srp1 | shown in (*85*) | no |
| Sla1 | Kap95, Srp1 | shown in (*86*) | no |
| Iwr1 | Kap95, Srp1 | shown in (*87*) | yes |
| Hxk2 | Kap95, Srp1 | shown in (*88*) | no |
| Rrp6 | Kap95, Srp1 | shown in (*89*) | no |
| Siz1 | Kap95, Srp1 | shown in (*90*) | no |
| Upf3 | Kap95, Srp1 | reviewed in (*36*) | no |
| Pcl5 | Kap95 | shown in (*91*) | no |
| Srp1 | Kap95 | shown in (*92*) | yes |
| Sts1 | Srp1 | shown in (*93*) | no |
| Gln3 | Srp1 | shown in (*94*) | no |
| Sto1 | Kap95, Srp1 | shown in (*95*, *96*) and reviewed in (*36*) | yes |
| Swi6 | Srp1 | reviewed in (*36*) | no |
| Clb2 | Kap95, Srp1 | shown in (*36*) | no |
| Gcn4 | Kap95, Srp1 | reviewed in (*36*) | no |
| Prp20 | Kap95, Srp1 | shown in (*36*) | no |
| Swi5 | Kap95, Srp1 | shown in (*36*) | no |
| Cdc6 | Kap95, Srp1 | shown in (*36*) | no |
| Prp8 | Kap95, Srp1 | shown in (*62*) | yes |

**Table S1:** True positive set of cargoes used for the analysis as depicted in **Fig. S3B**.

| Cargo | NLS (residues) | Reference | AF-Score (ipTM+pTM) |
|---|---|---|---|
| Nup1 | 1-123 | (*73*) | 0.65-0.73 |
| Nup2 | 1-50 | (*82*) | 0,76 |
| Ulp1 | 150-172 | (*57*) | 0.78-0,79 |
| Pct1 | 60-66 | (*84*) | 0.64-0.7 |
| Iwr1 | 9-43 | (*87*) | 0.6-0.7 |
| Sto1 | 2-30 | (*95*, *96*) | 0.8 |
| Prp8 | 96-117 | (*62*) | 0.7 |
| Opi1 | 109-112 | (*97*) | 0.33-0.56 |
| Heh2 | 124-137 | (*79*) | 0.67-0.74 |
| San1 | 180-200 | (*98*) | 0.66-0.76 |

**Table S2:** Literature reported cNLS for Srp1-Kap95 cargoes. A range of the AF-score is given when the NLS was contained in more than one sequence window. AF: AlphaFold.

**Table S3** is provided in a supplementary Excel sheet.

**Table S3:** Physical parameters and scores of protein-protein interfaces modeled by AlphaFold-Multimer. The columns show the following: “Model” – name of the target and region (in AA) that was modeled, “Interface residues” – number of residues at the interface, “Polar”, ”Hydrophobic”, and “Charged” – fractions of the corresponding type of residues at the interface, “Hydrogen bonds” – number of potential hydrogen bonds, “Salt bridges” – number of potential salt bridges, “Solvation free energy” - solvation free energy gain upon formation of the interface, “*P*-value” - the *P*-value of the observed solvation free energy gain (*P*-value < 0.5 implies higher likelihood that the interaction is specific (*99*),“PI score” – PI score of the interface, with values > 0 indicating that the interface resembles real interfaces. All values were calculated with PI_score pipeline (*100*) using CCP4 (*101*) and PISA (*99*) programs included in the pipeline. The models and crystal structures were minimized using GROMACS (*102*) prior to analysis. Cells with ipTM+pTM score color by value, from high to low, using color gradient from red to green. Blank cells mean that no interface was detected by PI_score.

| Strain Name | Genotype | Source |
|---|---|---|
| **BY4741 (wildtype)** | *MATa his3Δ1 leu2Δ0 met15Δ0, ura3Δ0* | provided by Patil lab |
| **Srp1-StrepII** | (BY4741)*srp1-strepII* | this study |
| **Kap95-StrepII** | (BY4741)*kap95-strepII* | this study |
| **Kap104-StrepII** | (BY4741)*kap104-strepII* | this study |
| **Kap114-StrepII** | (BY4741)*kap114-strepII* | this study |
| **Kap120-StrepII** | (BY4741)*kap120-strepII* | this study |
| **Kap121-StrepII** | (BY4741)*kap121-strepII* | this study |
| **Kap122-StrepII** | (BY4741)*kap122-strepII* | this study |
| **Kap123-StrepII** | (BY4741)*kap123-strepII* | this study |
| **Nmd5-StrepII** | (BY4741)*nmd5-strepII* | this study |
| **Sxm1-StrepII** | (BY4741)*sxm1-strepII* | this study |
| **Mtr10-StrepII** | (BY4741)*mtr10-strepII* | this study |
| **GFP** | pRS316 (tef1-promoter::*gfp*::cyc1-terminator::ura3) | this study |
| **Ino80(NLS)-GFP** | pRS316 (tef1-promoter:*ino80(1255-1374)-gfp*::cyc1-terminator::ura3) | this study |
| **Prp8(NLS)-GFP** | pRS316 (tef1-promoter::*prp8(502-621)-gfp*::cyc1-terminator::ura3) | this study |
| **Rps5(NLS)-GFP** | pRS316 (tef1-promoter::*rps5(238-393)-gfp*::cyc1-terminator::ura3) | this study |
| **Nup60(NLS)-GFP** | pRS316 (tef1-promoter::*nup60(1-159)-gfp*::cyc1-terminator::ura3) | this study |
| **Pop1(NLS)-GFP** | pRS316 (tef1-promoter::*pop1(316-444)-gfp*::cyc1-terminator::ura3) | this study |
| **Pct1(NLS-Srp1)-GFP** | pRS316 (tef1-promoter::*pct1(76-195)-gfp*::cyc1-terminator::ura3) | this study |
| **Pct1(NLS-Kap95)-GFP** | pRS316 (tef1-promoter::*pct1(247-366)-gfp*::cyc1-terminator::ura3) | This study |

**Table S4:** Yeast strains and corresponding genotypes used in this study.
